# *Rpl24^Bst^* mutation suppresses colorectal cancer by promoting eEF2 phosphorylation via eEF2K

**DOI:** 10.1101/2021.05.05.442715

**Authors:** John R. P. Knight, Nikola Vlahov, David M. Gay, Rachel A. Ridgway, William J. Faller, Christopher G. Proud, Giovanna R. Mallucci, Tobias von der Haar, C. Mark Smales, Anne E. Willis, Owen J. Sansom

## Abstract

Increased protein synthesis supports the rapid proliferation associated with cancer. The *Rpl24^Bst^* mutant mouse reduces the expression of the ribosomal protein RPL24 and has been used to suppress translation and limit tumorigenesis in multiple mouse models of cancer. Here we show that *Rpl24^Bst^* also suppresses tumorigenesis and proliferation in a model of colorectal cancer with two common patient mutations, *Apc* and *Kras*. In contrast to previous reports, *Rpl24^Bst^* mutation has no effect on ribosomal subunit abundance but suppresses translation elongation through phosphorylation of eEF2, reducing protein synthesis by 40% in tumour cells. Ablating eEF2 phosphorylation in *Rpl24^Bst^* mutant mice by inactivating its kinase, eEF2K, completely restores the rates of elongation and protein synthesis. Furthermore, eEF2K activity is required for the *Rpl24^Bst^* mutant to suppress tumorigenesis. This work demonstrates that elevation of eEF2 phosphorylation is an effective means to suppress colorectal tumorigenesis with two driver mutations. This positions translation elongation as a therapeutic target in colorectal cancer, as well as other cancers where the *Rpl24^Bst^* mutation has a tumour suppressive effect in mouse models.

## Introduction

Tumour cells require rapid protein synthesis to acquire sufficient biomass in order to divide and, as such, protein synthesis is directly regulated by many oncogenic signalling pathways (Proud 2019; Robichaud et al. 2019; Smith et al. 2021). As well as exploiting protein synthesis to drive cell division, cancers use translation to selectively synthesise a proteome geared towards proliferation, survival and immune evasion. For example, in colorectal cancer (CRC) translation of the mRNA encoding the proto-oncogene c-MYC is selectively upregulated by eIF4E and mTORC1 signalling (Knight et al. 2020a). Likewise, independent reports have shown that the expression of the immune suppressive ligand PD-L1 is maintained on tumour cells by the activity of the translation factors eIF4A and eIF5B as well as phosphorylation of eIF4E and eIF2 (Suresh et al. 2020; Xu et al. 2019; Cerezo et al. 2018).

*APC* is the most commonly mutated gene in CRC, followed by *TP53* and then *KRAS* (Guinney et al. 2015). We have previously shown that *Apc*-deficient mouse models of CRC are dependent on fast translation elongation, a process which can be suppressed by rapamycin leading to near complete reversal of tumorigenesis (Faller et al. 2015). This approach has had clinical success, where rapamycin (sirolimus) regressed *APC*-deficient polyps of familial adenomatous polyposis patients in two independent clinical trials (Yuksekkaya et al. 2016; Roos et al. 2020). Clinical data also suggest that CRCs increase translation elongation to potentiate proliferation, exemplified by the lower expression of the elongation inhibiting kinase eEF2K correlating with worse patient survival (Ng et al. 2019). However, the regulation of translation elongation in CRC is complex, notably being influenced by specific cancer-associated mutations. We have shown that mutation of *Kras* drives resistance to rapamycin in *Apc*-deficient models both in terms of its effect on elongation and on proliferation (Knight et al. 2020a). This is consistent with *KRAS*-mutant CRCs being resistant to rapalogues, and other therapeutics (Ng et al. 2013; Spindler et al. 2013; DeStefanis et al. 2019) and highlights the unmet need for effective therapies against *KRAS*-mutant cancers. Although a recently developed compound covalently targeting KRAS^G12C^ mutation has shown remarkable potency against this specific mutation (Hong et al. 2020).

Evidence suggests that targeting translation in *KRAS*-mutant colorectal cancer can still be effective. As well as our recent study re-sensitizing *Kras*-mutant CRCs to rapamycin by targeting translation initiation (Knight et al. 2020a), we have demonstrated that *Kras*-mutant models of CRC depend upon the transporter SLC7A5 to maintain protein synthesis by facilitating the influx of amino acids (Najumudeen et al. 2021). These data support protein synthesis as a tractable target in CRC, with the discovery of additional factors regulating these pathways only improving the potential to target protein synthesis in the clinic (Knight and Sansom 2021).

In this study we analyse the previously characterised *Rpl24^Bst^* mutation in models of CRC with *Apc*-deletion and *Kras* mutations. This spontaneously arising 4 nucleotide deletion in the *Rpl24* gene, which encodes RPL24 (a component of the 60S ribosomal subunit also called large ribosomal protein subunit eL24), disrupts splicing of its mRNA, effectively resulting in a *Rpl24* heterozygous animal (Oliver et al. 2004). Animals present with impaired dorsal pigmentation and malformed tails, among other defects, leading to the designation of a belly spot and tail (Bst) phenotype and the *Rpl24^Bst^* designation. This tool has been used to suppress overall protein synthesis in genetically engineered mouse models of c-MYC driven B-cell lymphoma (BCL), *Pten*-deficient T-cell acute lymphoblastic leukaemia (T-ALL) and T-cell specific *Akt2* activation (Barna et al. 2008; Signer et al. 2014; Hsieh et al. 2010). In these studies, tumorigenesis increased total protein synthesis, which was rescued by combination with the *Rpl24^Bst^* mutation. Suppression of protein synthesis was sufficient to slow tumorigenesis, with some *Rpl24^Bst/+^* mice surviving over 3 times longer than the median survival of tumour model mice wild-type for *Rpl24*. However, the means by which the *Rpl24^Bst^* mutation suppresses protein synthesis was not addressed in these studies, instead deferring to the original observation that there is likely a defect in ribosome production (Oliver et al. 2004).

Here we show that decreased expression of RPL24 suppresses proliferation and extends survival in *Apc*-deficient *Kras*-mutant pre-clinical mouse model of CRC. Importantly, we find that reduced RPL24 does not alter the available pool of ribosomal subunits, as previously suggested, but instead alters signalling that regulates a translation factor. Specifically, we observe increased phosphorylation of eEF2, an event that negatively regulates translation elongation. We directly measure translation elongation to show that the *Rpl24^Bst^* mutation suppresses protein synthesis at the elongation step, consistent with increased phosphorylation of eEF2. Relieving P-eEF2 by inactivating its inhibitory kinase, eEF2K, completely restores translation elongation and protein synthesis rates as well as reversing the beneficial effect of *Rpl24^Bst^* mutation in our tumour models. Interestingly, we find that the *Rpl24^Bst^* mutation has no effect in *Kras*-wild-type models. We attribute this to a specific requirement for physiological RPL24 in *Kras*-mutant cells, which may provide additional mechanisms to target these cells clinically.

Finally, we provide evidence from transcriptomic and proteomic analyses of patient tissue that supports the signalling pathways uncovered in our pre-clinical models being altered in the human disease. Altogether this work demonstrates that the *Rpl24^Bst^* mutation is tumour suppressive in a solid tumour model, and elucidates an unexpected mode of action underlying its impact on protein synthesis. This has implications for targeting translation elongation in cancer and provides mechanistic insight to supplement the previously published efficacy of the *Rpl24^Bst^* mouse in models of cancer and other diseases.

## Results

### RPL24^Bst^ mutation does not alter intestinal homeostasis but suppresses the rate of translation

Prior to addressing the role of RPL24 in intestinal tumorigenesis we first analysed whether the *Rpl24^Bst^* mutation had any effect on normal intestinal homeostasis (Figure 1A). We observed a reduction in RPL24 expression (Figure 1B), but no dramatic differences in intestinal architecture, crypt proliferation shown by BrdU incorporation (Figure 1A) or abundance of stem cells (*Olfm4*), Paneth cells (lysozyme) or goblet cells (AB/PAS) (Figure S1A). Similarly, homeostasis in the colons of *Rpl24^Bst^* mutant mice was unaffected, exemplified by no change in proliferation scored by BrdU incorporation (Figure S1C). In accordance with these *in vivo* observations, *ex vivo* organoids made from the small intestines of *Rpl24^Bst/+^* mice established in culture and grew comparably to wild-type controls (Figures 1C and S1D). Surprisingly, we measured a >40% reduction in total protein synthesis, by ^35^S-methionine labelling, in *Rpl24^Bst/+^* organoids compared to wild-type counterparts (Figure 1D). Therefore, despite no change in homeostasis, *Rpl24^Bst^* mutation had a dramatic effect on protein synthesis. This indicates a resistance to reduced protein synthesis in the wild-type intestine, which appear to function normally despite a dramatic reduction in protein output.

**Figure 1:**
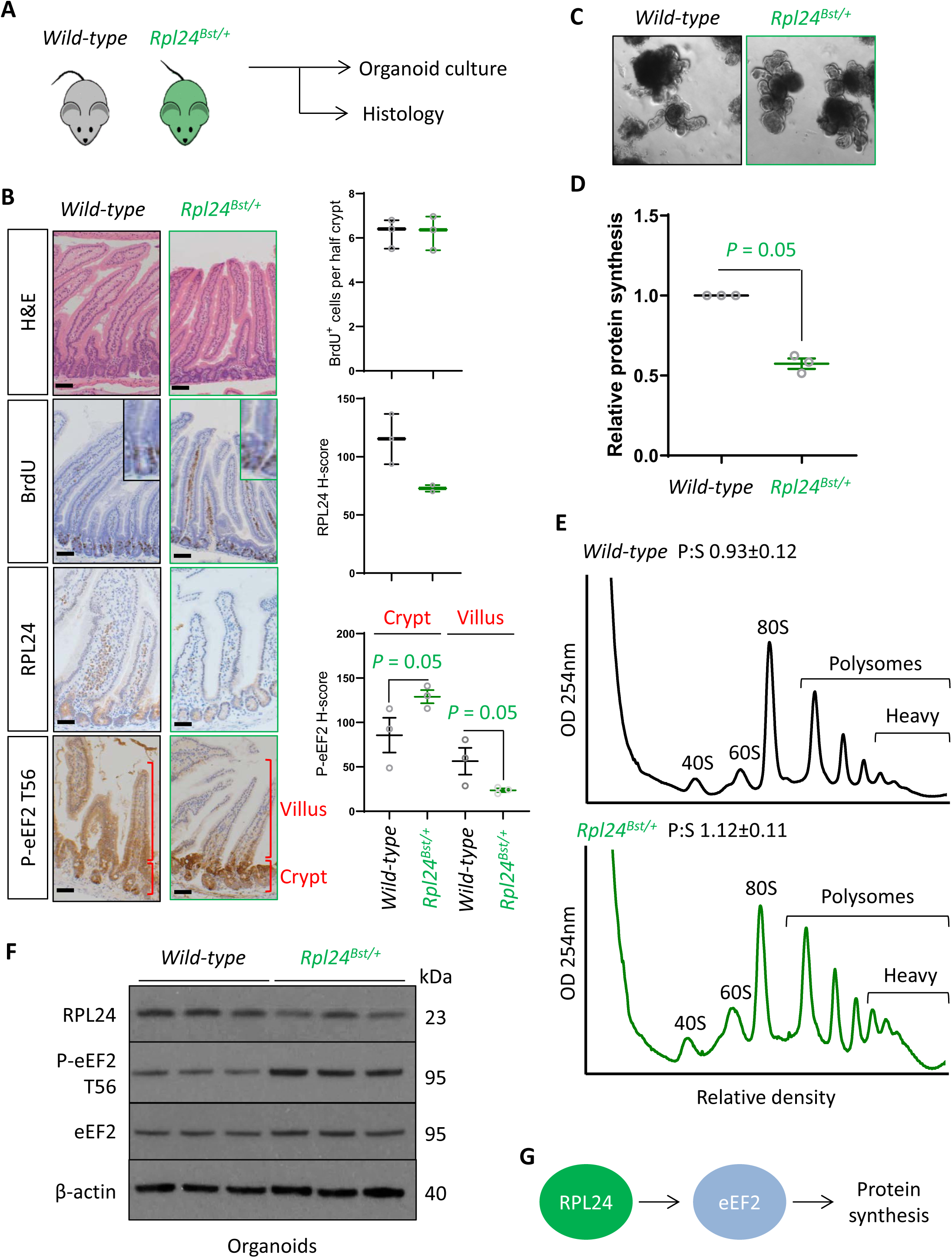
*Rpl24^Bst^* mutation slows translation elongation but does not affect homeostasis in the intestinal epithelium: (A) Schematic representation of experimental procedure. Intestines from wild-type or *Rpl24^Bst/+^* mice were analysed by histology or processed to make intestinal organoids. (B) Staining for H&E, BrdU, RPL24 andP- eEF2 T56 in sections from the small intestines of wild-type and *Rpl24^Bst/+^* mice. Red brackets in P-eEF2 staining indicates crypts and villi, correspondint to quantification to the right. Bars represent 50µm. Graphs on the right show scoring for BrdU positive cells, and H-score calculated for RPL24 and P-eEF2 T-56, plotted ±SEM. Significance was determined by one tailed Mann Whitney U test. (C) Micrographs of small intestinal organoids generated from wild-type or *Rpl24^Bst/+^* mice. (D) Protein synthesis rate quantified by ^35^S-methionine incorporation in wild-type or *Rpl24^Bst/+^* organoids (n=3), expressed relative to the wild-type protein synthesis rate (= 1). Data are from 3 biologically independent organoid lines for each genotype represented ±SEM with significance determine by Mann Whitney U test. (E) Representative polysome profiling from wild-type or *Rpl24^Bst/+^* organoids. Average polysome:sub-polysome ratios from 3 independent organoid lines per genotype are shown above each profile ±SEM. (F) Western blotting from protein lysates generated from 3 biologically independent organoid lines for each genotype. There is a 50% reduction in RPL24 and a 75% increase in P-eEF2 T56 in the *Rpl24^Bst/+^* organoids. (G) Schematic of the regulation of the potential role of RPL24 in regulating protein synthesis via eEF2.

We then investigated how *Rpl24^Bst^* mutation suppresses translation. Performing sucrose density gradients to quantify the number of ribosomes engaged in active translation we observed an increase in ribosomes bound to mRNAs in polysomes in *Rpl24^Bst/+^* organoids, particularly the heavy polysomes (Figure 1E). This appears to contradict the reduction in global protein synthesis observed in Figure 1D. However, ourselves and others have previously observed increased polysomes in conjunction with reduced protein synthesis in model systems where translation elongation is reduced (Knight et al. 2015; Faller et al. 2015). In these instances slowed translation elongation increased the abundance of polysomes via changes in signalling to the elongation factor eEF2. We therefore assayed the regulatory phosphorylation of eEF2 (threonine 56 / T56) in wild-type and *Rpl24^Bst^* mutant samples. This phosphorylation event excludes eEF2 from the ribosome thereby impairing the translocation step of translation elongation, reducing protein synthesis (Ryazanov and Davydova 1989; Carlberg et al. 1990).

In small intestinal tissue assayed by IHC we observed an increase in the P-eEF2 T56, specifically in the proliferating crypt and transit amplifying zone of *Rpl24^Bst/+^* mouse intestines (Figure 1B). Likewise we observed a 75% increase in P-eEF2 T56 in lysates generated from *Rpl24^Bst/+^* organoids compared to wild-type organoids (Figure 1D). These organoids also showed a 50% reduction in RPL24 expression by western blot (Figure 1D). Thus, in these two proliferative settings (intestinal crypts *in situ* and *ex vivo* organoids) *Rpl24^Bst^* mutation increases the phosphorylation of eEF2, which is known to suppress translation. Surprisingly, we observed that in the differentiated villus, *Rpl24^Bst^* mutation suppressed P-eEF2 T56 (Figure 1B). The reasons for this are unclear, but may relate to different cell functions in the two compartments. This effect on P-eEF2 T56 is specific, as we observed no effect on the phosphorylation of other translation related proteins, 4E-BP1 or RPS6 (at S240/S244), readouts for modulation of signalling downstream of mTORC1 (Figure S1A-B).

Altogether, this analysis of wild-type tissue indicates that physiological RPL24 expression is not required for proliferation or function, but reduction of RPL24 expression reduces the rate of translation. Interestingly, this involves regulation of translation elongation and correlates with increased P-eEF2 (Figure 1G). Previously the *Rpl24^Bst^* mutation had been suggested to suppress ribosome biogenesis, resulting in uneven 40S and 60S ratios. However, we observed no alteration in the relative levels of the 40S and 60S subunits in sucrose density gradients (Figure S1E), consistent with previous reports that RPL24 deletion has little effect on ribosome biogenesis (Barkić et al. 2009).

The wild-type mouse intestine regenerates following γ-irradiation, dependent on Wnt and MAPK signalling pathways. These pathways are often deregulated in colorectal tumours, such that this intestinal regeneration acts as a surrogate for oncogenic potential, with reduced regenerative capacity indicative of reduced tumorigenic proliferation (Faller et al. 2015). We observe that mutation of *Rpl24* restricts regeneration of the small intestine (Figure S1F). Thus, RPL24 expression enables regeneration, which may correlate with effects in tumorigenesis.

### RPL24 is required for proliferation in Apc-deficient Kras-mutant intestinal tumours

Next, we analysed the effect of the *Rpl24^Bst^* mutation on a model of CRC driven by tamoxifen-inducible *Villin^CreER^* mediated deletion of *Apc* and activation of KRAS with a G12D mutation. This model can be used with homozygous deletion of *Apc* (*Apc^fl/fl^*) where intestinal hyperproliferation generates a short term (3-4 days) hyperproliferation model or with heterozygous deletion of *Apc* (*Apc^fl/+^*) where intestinal adenomas form following spontaneous loss of the second copy of *Apc*. The recombination of a lox-STOP-lox allele at the endogenous *Kras* locus expresses a constitutively active G12D mutant form of the protein. Mice were generated with the *Apc^fl/fl^* and *Kras^G12D^* alleles with and without the *Rpl24^Bst^* mutation (Figure 2A). Hyperproliferation in the *Apc^fl/fl^ Kras^G12D/+^ Rpl24^Bst/+^* mutant mice was significantly suppressed compared to *Apc^fl/fl^ Kras^G12D/+^* control mice (Figure 2B and C). Reduced RPL24 expression was confirmed by IHC (Figure 2C) and coincided with increased P-eEF2 throughout the proliferative crypt area of the *Apc^fl/fl^ Kras^G12D/+^ Rpl24^Bst/+^* intestine (Figure 2C and S2A). Hyper-proliferation in the colon mirrored that of the small intestine, with reduced proliferation in mice mutant for *Rpl24* compared to those expressing wild-types levels of RPL24 (Figure S2B). In parallel, organoids were derived from the small intestines of the same genotypes (*Apc^fl/fl^ Kras^G12D/+^* and *Apc^fl/fl^ Kras^G12D/+^ Rpl24^Bst/+^*) and their growth compared. *Rpl24^Bst^* mutation resulted in significantly less proliferation *ex vivo* (Figure 2D), consistent with the *in vivo* experiment shown in Figure 2B.

**Figure 2:**
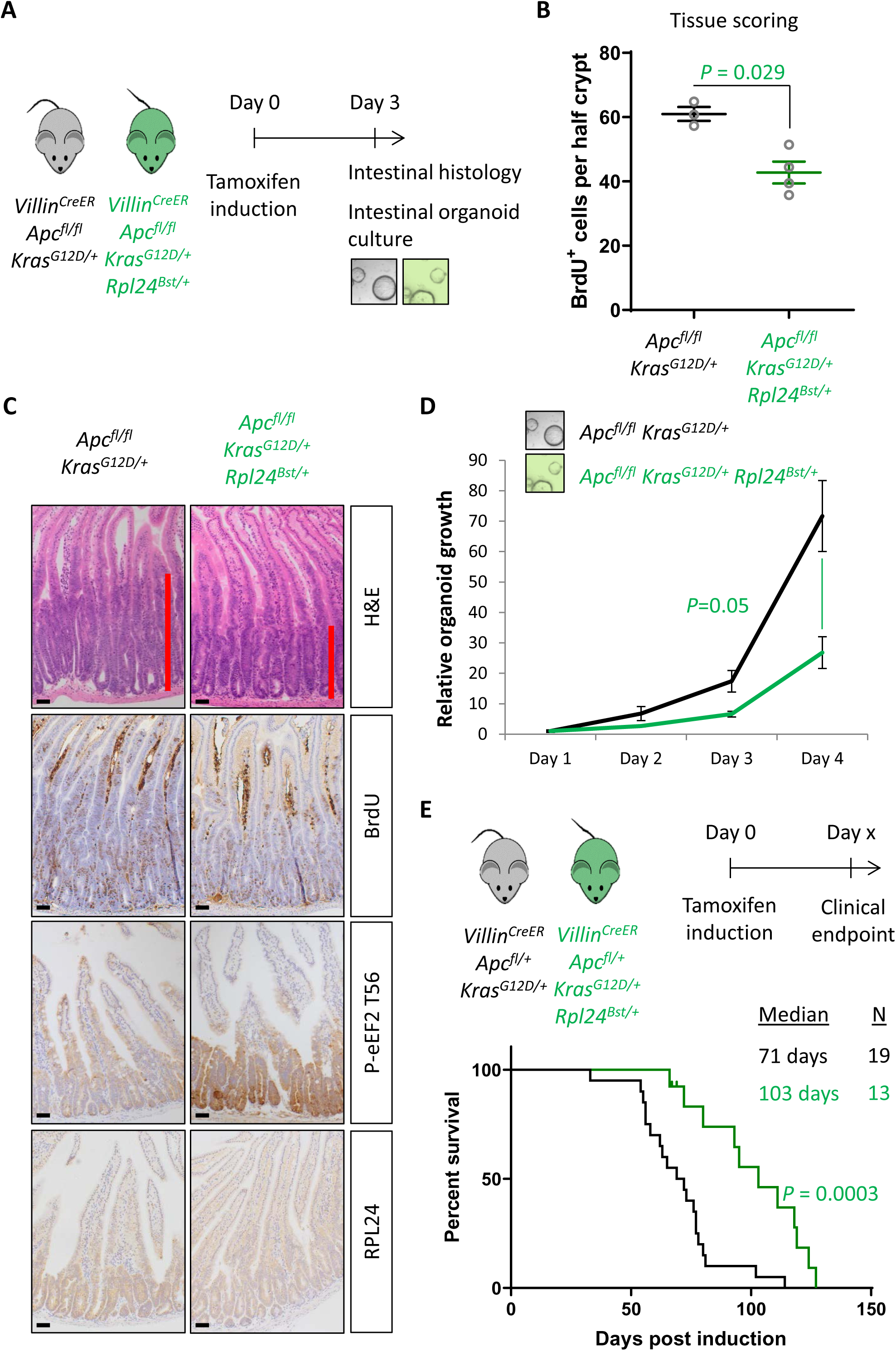
*Rpl24^Bst^* mutation suppresses proliferation and extends survival in an *Apc*-deficient *Kras*-mutant mouse model of CRC: (A) Schematic representation of experimental protocols. *Villin^CreER^ Apc^fl/fl^ Kras^G12D/+^* or *Villin^CreER^ Apc^fl/fl^ Kras^G12D/+^ Rpl24^Bst/+^* mice were induced by intraperitoneal injection of tamoxifen at 80mg/kg then intestinal tissue analysed 3 days later. Tissue was taken for histological analysis or processed into intestinal organoids. (B) Quantification of BrdU incorporation in small intestinal crypts following deletion of *Apc* and activation of *Kras*, with (n=3) or without (n=4) *Rpl24^Bst^* mutation. Data are represented as the mean number of BrdU positive cells per half crypt from >20 crypts per mouse, ±SEM. Significance was determined by Mann Whitney U test. (C) Representative images of intestines from the same experiment as in (B), stained for H&E, BrdU, P-eEF2 T56 and RPL24. The red bar on the H&E images indicates the extent of the proliferative zone. Bars represent 50µm. (D) *Apc^fl/fl^ Kras^G12D/+^* organoids with or without *Rpl24^Bst^* mutation were grown for 4 days and growth relative to day 1 determined by Cell Titer Blue assay. Data show the mean ±SEM of n=3 independent organoid lines. Significance was determined by one tailed Mann Whitney U test. (E) Top: schematic of experimental protocol. *Villin^CreER^ Apc^fl/+^ Kras^G12D/+^* or *Villin^CreER^ Apc^fl/+^ Kras^G12D/+^ Rpl24^Bst/+^* mice induced with 80mg/kg tamoxifen then monitored until clinical endpoint. Survival plot for these genotypes for the days post induction that they reached endpoint. The median survival and n number for each cohort is shown and significance determined by Mantel-Cox test. Censored subjects were removed from the study due to non-intestinal phenotypes.

In the tumour model, where a single copy of *Apc* is deleted, *Apc^fll+^ Kras^G12D/+^ Rpl24^Bst/+^* mice lived on average 32 days longer than *Apc^fll+^ Kras^G12D/+^* controls, an extension of survival of 45% (Figure 2E). *Apc^fll+^ Kras^G12D/+^ Rpl24^Bst/+^* organoids derived from the adenomas in this tumour model grew more slowly than controls (Figure S2C). There was no significant difference in the number of tumours at experimental endpoint (Figure S2D), indicating that adenomas can form but take longer to reach a clinically significant burden. Therefore, RPL24 enables proliferation in *Apc*-deficient, KRAS-activated cells within the intestinal epithelium of the mouse.

### RPL24 maintains translation elongation in Apc-deficient Kras-mutant intestinal tumour models

The suppression of tumorigenesis in the *Apc*-deficient *Kras*-mutant model correlated with increased phosphorylation of eEF2 (Figure 2C and S2A). To investigate this further we used 3 methods to measure the rate of translation; polysome profiling, ^35^S-methionine labelling and harringtonine run-off assays (Figure 3A). Polysome profiling from extracted crypts from *Apc^fl/fl^ Kras^G12D/+^* and *Apc^fl/fl^ Kras^G12D/+^ Rpl24^Bst/+^* showed an increase in polysomes with the *Rpl24* mutation (Figure 3B), and notably a significant increase in the quantity of heavy polysomes (Figure 3C). Intestinal organoids of the same genotype showed a 35% reduction in ^35^S-methionine incorporation (Figure 3D). These same organoids had a greater than 40% decrease in elongation rate measured by harringtonine run-off (Figures 3D and S3A). Together these data provide strong evidence that normal RPL24 expression is required to maintain translation elongation in this CRC model. Polysome profiles and protein synthesis rate measurements from *Apc^fl/+^ Kras^G12D/+^ Rpl24^Bst/+^* adenoma cultures also showed more polysomes and lower protein synthesis compared to control adenoma cultures (Figures S3B and C). Furthermore, the reduced protein synthesis rate does not correlate with differences in free ribosomal subunit availability as the ratio of 40S to 60S subunits is unchanged by the *Rpl24* mutation in *Apc*-deficient *Kras*-mutant cells (Figure S3D).

**Figure 3:**
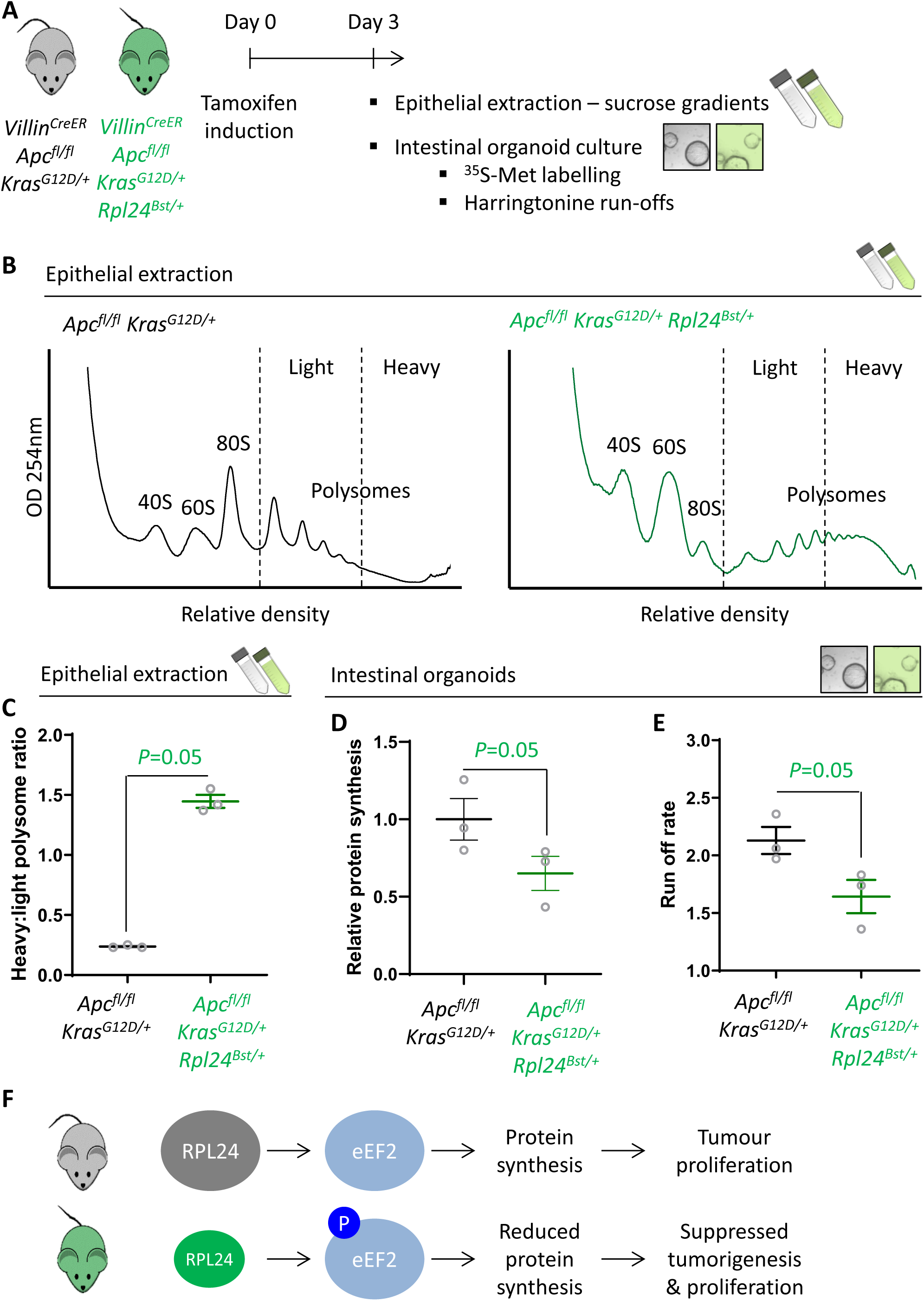
*Rpl24^Bst^* mutation slows translation elongation in *Apc*-deficient *Kras*-mutant mouse models of CRC: (A) Schematic representation of experimental approach. *Villin^CreER^ Apc^fl/fl^ Kras^G12D/+^* or *Villin^CreER^ Apc^fl/fl^ Kras^G12D/+^ Rpl24^Bst/+^* mice were induced by intraperitoneal injection of tamoxifen at 80mg/kg then intestinal tissue analysed 3 days later. Intestines were enriched for crypt epithelium for sucrose density analysis or processed into intestinal organoids. (B) Representative sucrose density polysome profiles generated from *Apc^fl/fl^ Kras^G12D/+^* intestinal extracts with or without the *Rpl24^Bst^* mutation. Subpolysomal components (40S, 60S and 80S) and polysomes are labelled, with the polysomes also split pictorially into light and heavy. (C) Quantification of the heavy:light polysome ratio from the experiment in (B). Data show the mean of analysis from 3 mice ±SEM with significance determined by one tailed Mann Whitney U test. (D) Relative protein synthesis rate quantified by ^35^S-methionine incorporation in *Apc^fl/fl^ Kras^G12D/+^* three biologically independent organoid lines either wild-type or mutant for *Rpl24^Bst^*. Data are represented ±SEM with significance determine by Mann Whitney U test. (E) Ribosome run-off rate determined in *Apc^fl/fl^ Kras^G12D/+^* small intestinal organoid lines either wild-type or mutant for *Rpl24^Bst^* (n=3 per genotype). Data are represented as the mean of 3 biological replicates ±SEM with significance determine by Mann Whitney U test. Raw data are available in Figure S3A. (F) Schematics of the regulation of protein synthesis and tumour proliferation downstream of RPL24. Smaller RPL24 in bottom scheme represents reduced RPL24 expression. ‘P’ represents phosphorylation of eEF2.

Interpreting this molecular analysis in conjunction with the effects on tumorigenesis leads to the conclusion that RPL24 expression maintains translation elongation and protein synthesis rates, which in turn maintain tumour-related proliferation (Figure 3F). Furthermore, suppressing RPL24 expression increases P-eEF2, which decreases the rate of elongation and overall protein synthesis and correlates with suppressed tumorigenesis and proliferation.

### Rpl24 mutation has no effect in CRC models expressing wild-type Kras

In parallel to analysing the effect of *Rpl24* mutation in *Apc*-deficient *Kras*-mutant intestinal tumours, we also assessed its role in *Apc*-deficient models wild-type for *Kras*. We have previously shown that these are dependent on signalling from mTORC1 to maintain low levels of P-eEF2, and that this can be suppressed by rapamycin treatment to great therapeutic benefit (Faller et al. 2015). We observed that the hyperproliferation in the small intestine or colon of *Apc^fl/fl^* was not reduced by the *Rpl24^Bst^* mutation (Figure 4A and B, and S4A). Furthermore, in both germline *Apc^Minl+^* and inducible *Apc^fll+^* models of *Apc*-deficiency we see no benefit of the *Rpl24^Bst^* mutation, with no difference in survival or tumour development (Figures 4C and S4B). In agreement, *Apc^fl/fl^* organoids with the *Rpl24^Bst^* mutation grew at an identical rate to those wild-type for *Rpl24* in culture (Figure 4D). Together these data demonstrate that reduced RPL24 expression does not limit tumorigenesis in *Apc*-deficient CRC models with wild-type *Kras*.

**Figure 4:**
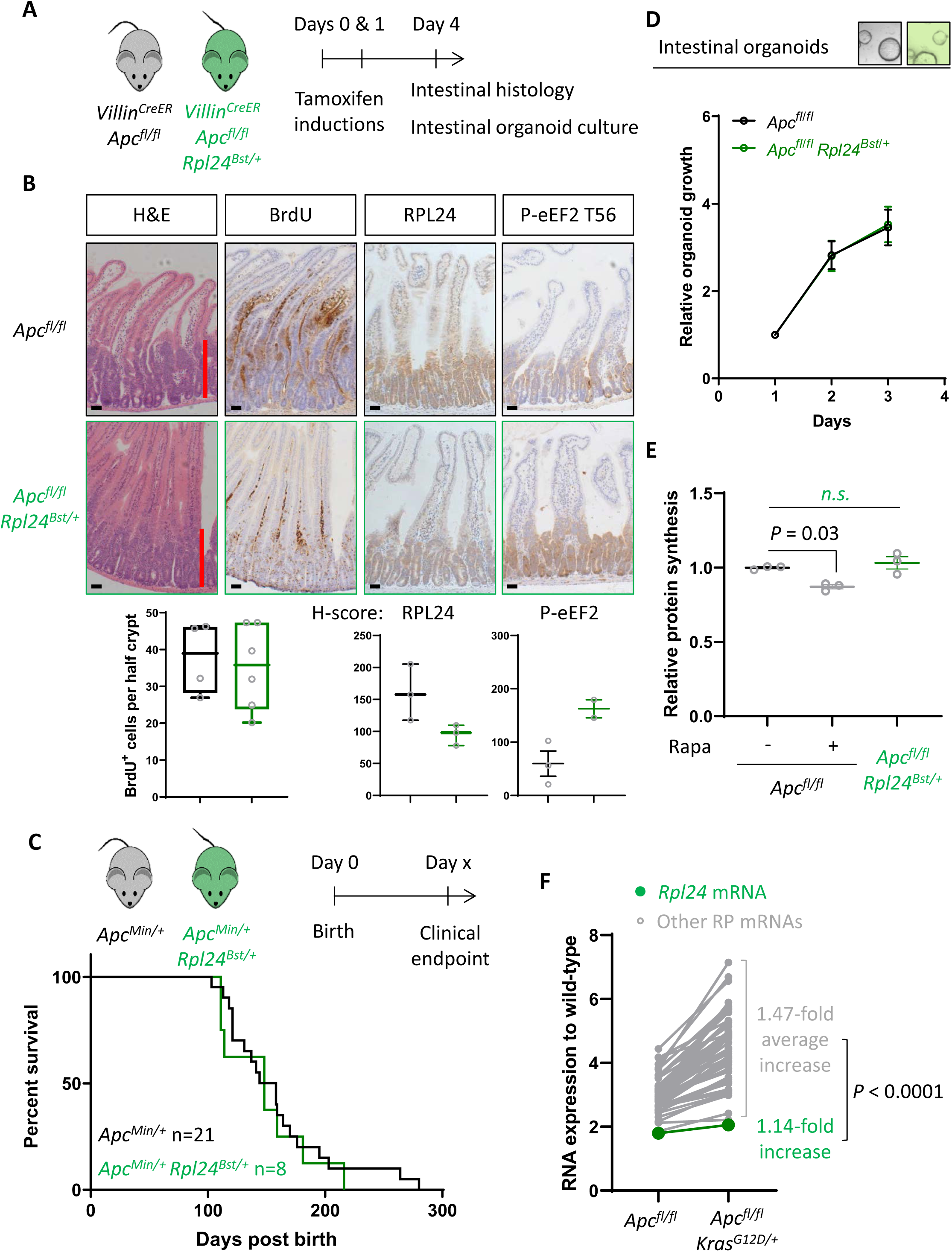
*Rpl24^Bst^* mutation does not suppress proteins synthesis or proliferation in *Apc*-deficient *Kras* wild-type mouse models of CRC: (A) Schematic representation of experimental approach. *Villin^CreER^ Apc^fl/fl^* or *Villin^CreER^ Apc^fl/fl^ Rpl24^Bst/+^* mice were induced by 2 intraperitoneal injection of tamoxifen at 80mg/kg on days 0 and 1 then intestinal tissue analysed on day 4. Intestines analysed histologically or intestinal organoids generated. (B) Top: representative micrographs showing proliferation as BrdU positivity and extent of proliferation as a red bar in H&E image. RPL24 and P-eEF2 T56, staining is also shown for each genotype. Bars represent 50µm. Below: BrdU scoring from *Apc^fl/fl^* or *Apc^fl/fl^ Rpl24^Bst/+^* mouse intestines and H-scores for RPL24 and P-eEF2 T56 protein levels. For BrdU scoring BrdU was administered 2 h before sampling and at least 20 half crypts were scored per animal and the mean plotted ±SEM (C) *Apc^Min/+^* tumour model survival curve, for mice with and without *Rpl24^Bst^* mutation. Lack of a significant difference was determined by Mantel-Cox test. (D) Relative growth of *Apc^fl/fl^* and *Apc^fl/fl^ Rpl24^Bst/+^* small intestinal organoids over 3 days, measure by Cell-Titer blue assay. The average change in proliferation is plotted from 3 independent biological replicates per genotype. (E) Relative protein synthesis rates quantified from ^35^S-methionine incorporation into *Apc^fl/fl^*, *Apc^fl/fl^* treated with 250nM rapamycin for 24 h and *Apc^fl/fl^ Rpl24^Bst/+^* small intestinal organoids. Significant changes were calculated by one-way ANOVA analysis with Tukey’s multiple comparison. N=3 per genotype with the mean protein synthesis rate for each genotype plotted ±SEM. (F) Relative expression of ribosomal protein mRNAs in *Villin^CreER^ Apc^fl/fl^* and *Villin^CreER^ Apc^fl/fl^ Kras^G12D/+^* whole intestine samples, where wild-type tissue has been normalised to 1. The fold increase in expression from *Apc^fl/fl^* to *Apc^fl/fl^ Kras^G12D/+^* samples for *Rpl24* and the average of all other RP mRNAs is shown. Statistical analysis was by one sample t test of the other RP mRNA fold changes using the fold-change for *Rpl24* mRNA as the hypothetical mean.

Despite no effect on proliferation, we observed an increase in P-eEF2 in *Apc^fl/fl^ Rpl24^Bst/+^* intestines compared to *Apc^fl/fl^* (Figure 4B), showing that P-eEF2 is consistently increased in the intestinal crypts of *Rpl24^Bst/+^* mice. However, we see no change in the ratio of polysome to sub-polysomes in *Apc^fl/fl^ Rpl24^Bst/+^* intestines compared to *Apc^fl/fl^* (Figure S4C-D) and no change in protein synthesis rate between organoids of these same genotypes (Figure 4D). In contrast, rapamycin treatment significantly reduces protein synthesis in *Apc^fl/fl^* organoids treated in parallel (Figure 4D). From these data we conclude that the change in P-eEF2 does not limit the rate of protein synthesis which likely allows efficient tumorigenesis in these *Kras*-wild-type models of CRC. Importantly, neither P-4E-BP1 or P-RPS6 S240/244 are reduced by *Rpl24^Bst^* mutation in the *Apc^fl/fl^* model (Figure S4E), indicating that translation-promoting mTORC1 signalling remains high.

We hypothesised that the reason for the KRAS-specificity seen with the *Rpl24^Bst^* mutation may relate to expression levels between the different genotypes analysed. Using unbiased RNA sequencing data from wild-type, *Apc^fl/fl^* and *Apc^fl/+^ Kras^G12D/+^* small intestinal tissue we observed a consistent increase in ribosomal protein expression following *Apc* deletion, then again following KRAS activation (Figure S4F). This is consistent with previous reports (Smit et al. 2020), and a requirement to increase protein synthesis as a direct consequence of KRAS activation. Indeed, the mRNA for all ribosomal proteins with sufficient reads were increased on average nearly 1.5-fold by KRAS activation in the small intestine (Figure 4F). In contrast, the *Rpl24* mRNA was only increased by 1.14-fold following Kras mutation, despite a nearly two-fold increase following deletion of *Apc* (Figure 4F). This manifests as a significant difference in the RNA expression of *Rpl24* compared to the other ribosomal proteins. Therefore, RPL24 expression may be sufficient in *Rpl24^Bst^* mice in *Apc*-deleted models, but then becomes limiting following *Kras*-mutation due to the relatively limited upregulation of *Rpl24* expression accompanying KRAS activation.

### Genetic inactivation of eEF2K completely reverses the anti-proliferative benefit of Rpl24 mutation

Thus far we have demonstrated a correlation between the increase in P-eEF2 and the slowing of translation elongation following *Rpl24^Bst^* mutation. To test whether the slowing of elongation caused by *Rpl24^Bst^* mutation was dependent on P-eEF2, we used a whole-body point mutant of eEF2K, the kinase that phosphorylates eEF2, which almost completely inactivates its kinase activity (Gildish et al. 2012). We crossed this *Eef2k^D273A/D273A^* allele to the *Apc^fl/fl^ Kras^G12D/+^ Rpl24^Bst/+^* mice to generate *Apc^fl/fl^ Kras^G12D/+^ Rpl24^Bst/+^ Eef2k^D273A/D273A^* mice. In the short term hyperproliferation model, the inactivation of *Eef2k* completely reversed the suppression of proliferation seen in *Apc^fl/fl^ Kras^G12D/+^ Rpl24^Bst/+^* small intestines (Figure 5A-C), and the medial colon (Figure S5A). The kinase inactive *Eef2k* allele resulted in no detectable P-eEF2 (Figures 5C and S2A), and we previously reported no difference in hyper-proliferation between *Apc^fl/fl^ Kras^G12D/+^* and *Apc^fl/fl^ Kras^G12D/+^ Eef2k^D273A/D273A^* models (Knight et al. 2020a). The reversal in proliferation rate with *Eef2k* and *Rpl24^Bst^* mutations was also seen in intestinal organoid growth after 3 days (Figure 4D). Furthermore, inactivation of eEF2K reverted the survival benefit of the *Rpl24^Bst^* mutation in the *Apc^fll+^ Kras^G12D/+^* tumour model (Figure 4E). This experiment also shows the lack of effect of the *Eef2k^D273A/D273A^* mutation on tumorigenesis. Indeed, *Eef2k^D273A/D273A^* had no impact on tumorigenesis in inducible *Apc^fll+^* and germline *Apc^Minl+^* models of *Apc*-deficient intestinal tumorigenesis (i.e. expressing wild-type *Kras*) (Figures S5B-C). Heterozygous inactivation of one copy of *Eef2k* resulted in a slight reversal of the extension of survival associated with *Rpl24^Bst^* mutation in the *Apc^fll+^ Kras^G12D/+^* tumour model (Figure S5D).

**Figure 5:**
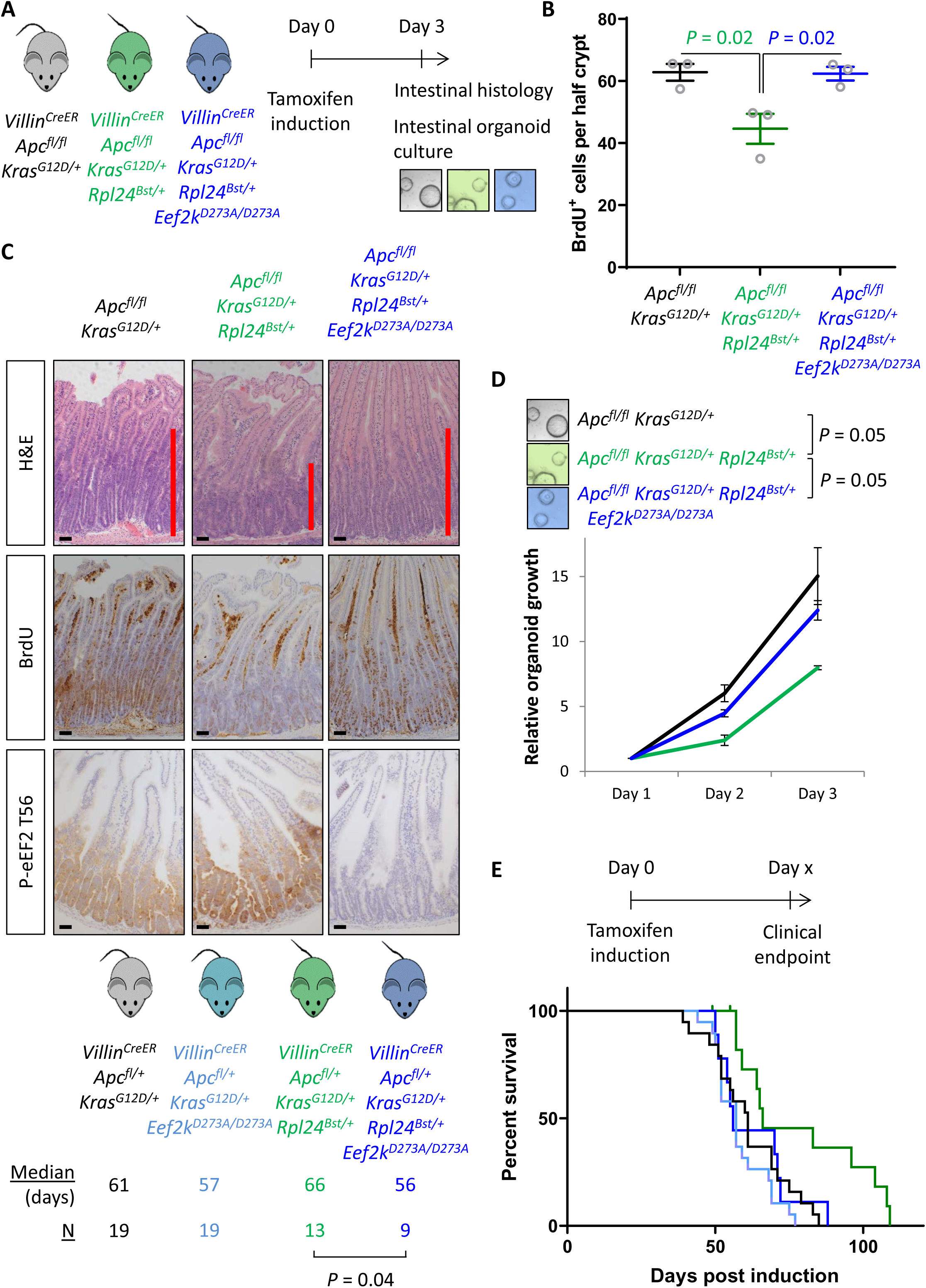
Genetic inactivation of eEF2K reverses the reduced tumorigenesis following *Rpl24^Bst^* mutation in *Apc*-deficient *Kras*-mutant models of CRC: (A) Schematic representation of experimental approach. *Villin^CreER^ Apc^fl/fl^ Kras^G12D/+^, Villin^CreER^ Apc^fl/fl^ Kras^G12D/+^ Rpl24^Bst/+^* or *Villin^CreER^ Apc^fl/fl^ Kras^G12D/+^ Rpl24^Bst/+^ Eef2k^D273A/D273A^* mice were induced by intraperitoneal injection of tamoxifen at 80mg/kg then intestinal tissue analysed 3 days later. Intestines were analysed histologically or processed into intestinal organoids. (B) BrdU incorporation quantified from within small intestinal crypts following deletion of *Apc* and activation of *Kras*, either wild-type of mutant for *Rpl24*, or mutant for *Rpl24* and *Eef2k*. Data are represented as the mean of at least 20 crypts per mouse ±SEM with significance determined by one-way ANOVA analysis with Tukey’s multiple comparison. N=3 per genotype. (C) Representative images of H&E, BrdU and P-eEF2 T56 staining of intestines from the same experiment as (B). Red bar on H&E indicates extent of proliferative zone. Bars represent 50µm. (D) Organoids deficient for *Apc* and with activated *Kras* with or without *Rpl24^Bst^* mutation, or mutant for both *Rpl24* and *Eef2k* were grown for 3 days and growth relative to day 1 determined by Cell Titer Blue assay. Data show the mean ±SEM of n=3 biologically independent organoid lines. Significance was determined by one tailed Mann Whitney U test. (E) Survival plot for *Apc Kras* ageing mice with or without the *Rpl24^Bst^* mutation, *Eef2K* mutation and with both *Rpl24* and *Eef2k* mutations. Median survival and n numbers for each cohort are shown and significance determined by Mantel-Cox test. Censored subjects were removed from the study due to non-intestinal phenotypes.

Therefore, mutation of *Rpl24* requires functional eEF2K to suppress proliferation and extend survival in this model of CRC. There was no alteration in the number of tumours in the ageing model at endpoint (Figure S4E), again identifying tumour cell proliferation, rather than tumour initiation, as the important factor regulated by RPL24 and eEF2K. These data also identify the effect of *Rpl24^Bst^* mutation on the tumour phenotype was entirely dependent upon eEF2K activity.

### Rpl24^Bst^ mutation suppresses translation exclusively via eEF2K/P-eEF2

Next, we addressed the molecular consequences of inactivation of eEF2K downstream of *Rpl24^Bst^* mutation. The reduction in protein synthesis that results from *Rpl24^Bst^* mutation is completely reversed by eEF2K inactivation (Figure 6A), with *Apc^fl/fl^ Kras^G12D/+^ Rpl24^Bst/+^ Eef2k^D273A/D273A^* organoids having an almost identical translation capacity as *Apc^fl/fl^ Kras^G12D/+^* controls. Similarly, the rate of ribosome run-off was also reverted in *Apc^fl/fl^ Kras^G12D/+^ Rpl24^Bst/+^ Eef2k^D273A/D273A^* compared to *Apc^fl/fl^ Kras^G12D/+^ Rpl24^Bst/+^* organoids, again to the same rate as controls with wild-type *Rpl24* (Figures 6B and S6A). In agreement, crypt fractions from *Apc^fl/fl^ Kras^G12D/+^ Rpl24^Bst/+^ Eef2k^D273A/D273A^* mice have a reduced number of heavy polysomes compared to *Apc^fl/fl^ Kras^G12D/+^ Rpl24^Bst/+^* crypt cells (Figure S6B), indicating faster translation elongation following eEF2K inactivation.

**Figure 6:**
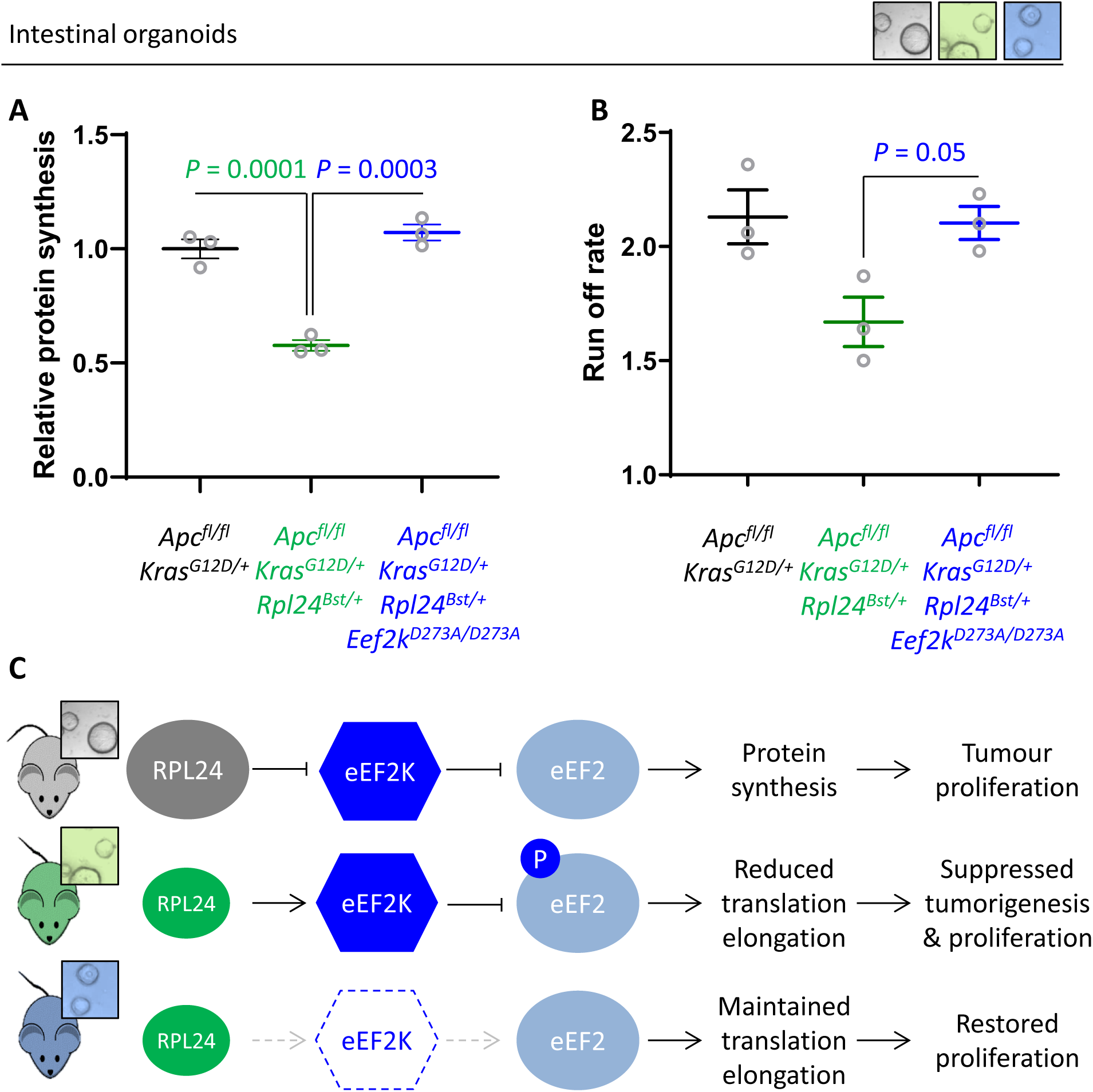
Genetic inactivation of eEF2K restores translation rates following *Rpl24^Bst^* mutation: (A) ^35^S-methionine incorporation to determine relative protein synthesis by in *Apc^fl/fl^ Kras^G12D/+^* small intestinal organoids wild-type or mutant for *Rpl24* or with both *Rpl24* and *Eef2k* mutations. Data are represented ±SEM with significance determined by one-way ANOVA analysis with Tukey’s multiple comparison. N=3 per genotype, each representing an independent organoid line. (B) Ribosome run-off rate determined in *Apc^fl/fl^ Kras^G12D/+^* small intestinal organoids mutant or wild-type for *Rpl24* or with both *Rpl24* and *Eef2k* mutations. Data are the mean of 3 biologically independent organoid lines represented ±SEM with significance determined by Mann Whitney U test. Raw data are available in Figure S5A. The run-off rate for *Apc^fl/fl^ Kras^G12D/+^* control organoids is reproduced from Figure 3E. (C) Schematic representation of findings in *Apc*-deficient *Kras-*mutant mouse and organoid models. Top: RPL24 expression maintains translation and proliferation by suppressing the phosphorylation of eEF2 by limiting eEF2K activity. Middle: reduced expression of RPL24 activates eEF2K, increasing P-eEF2, reducing translation elongation and suppressing tumorigenesis and proliferation. Bottom: inactivation of eEF2K reverts the phenotype in *Rpl24^Bst^* cells, due to the inability to phosphorylate and suppress eEF2. Elevated elongation rates correlate with increased proliferation following inactivation of eEF2K.

This has important implications for the function of RPL24, showing that ribosomes in *Rpl24^Bst^* mutant cells can elongate efficiently despite the reduction in RPL24 expression. However, reduced RPL24 increases P-eEF2 which inhibits elongation, with an absolute requirement for eEF2K for this (Figure 6C). Importantly, the combination of the *Rpl24* and *eEF2K* mutants shows that when P-eEF2 is abolished elongation occurs at normal speed, despite reduced expression of RPL24.

### The expression of RPL24, EEF2K and EEF2 is indicative of fast elongation in human CRC

Using pre-clinical mouse models we have demonstrated that physiological RPL24 expression maintains low eEF2K-mediated phosphorylation of eEF2 (Figure 7A). Hypo-phosphorylated eEF2 then ensures rapid protein synthesis enabling tumour proliferation *in vivo*. This requirement for RPL24 is specific to mutant-*Kras* models, indicating a likely relationship between KRAS and RPL24 that is beyond the scope of this work. We next sought to position this pre-clinical work in the context of clinical studies of the human disease. Using publicly available datasets for RNA expression in normal and colon cancer tissue we observe increased *RPL24* and *EEF2* expression in conjunction with reduced *EEF2K* expression (Figure 7B). This mirrors the signalling pathways in our mouse models which focus on ensuring high expression of active eEF2. Elevated *EEF2* message and reduced *EEF2K* message in these clinical samples is consistent with conservation of this signalling in the clinic. Similar results are seen for the protein expression of RPL24, eEF2K and eEF2 from colon adenocarcinoma samples (Figure S7A), and for the three mRNAs in rectal adenocarcinoma (Figure S7B). These expression analyses highlight the conservation of the proliferative tumour-associated signalling pathways characterised in our pre-clinical mouse models and patient samples from the clinic.

**Figure 7:**
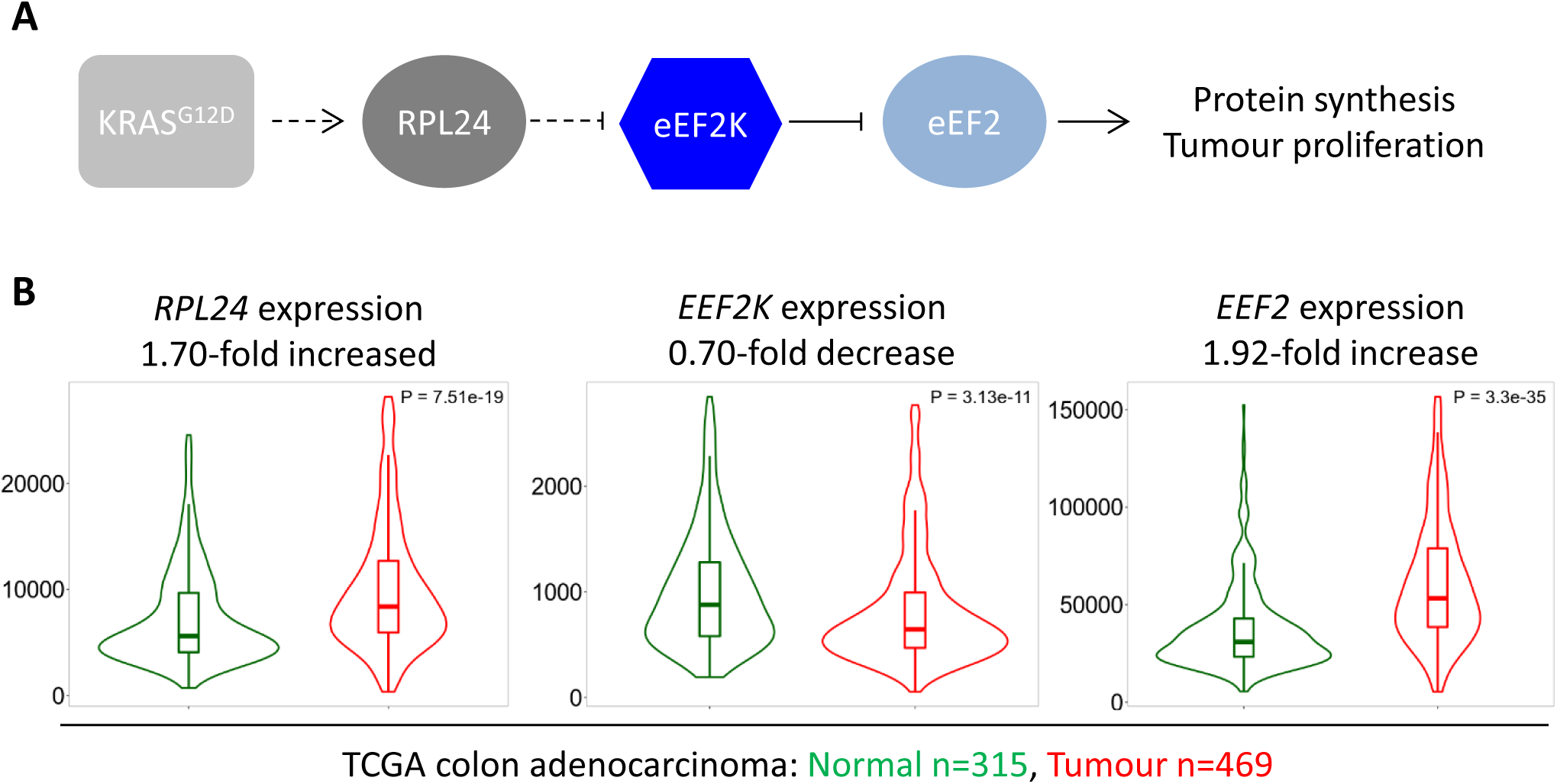
Expression of *RPL24*, *EEF2K* and *EEF2* is consistent with increased eEF2 activity in CRC tumours (A) Schematic of the findings presented here from pre-clinical mouse models. KRAS activation requires RPL24 expression to maintain low eEF2 phosphorylation. This occurs via a double negative regulation of eEF2K, whereby RPL24 suppresses eEF2K, which suppresses eEF2. eEF2 activity correlates with protein synthesis and proliferation rates. Dashed lines indicate indirect or undefined regulatory pathways. (B) RNA expression levels of RPL24, EEF2K and EEF2 between normal colon and colon adenocarcinoma samples using data extracted from The Cancer Genome Atlas by TNMplot. Relative expression changes are annotated, as well as *P* values for each transcript.

## Discussion

The original characterisation of the *Rpl24^Bst^* mutation identified a defect in ribosome biogenesis affecting the synthesis of 60S subunits (Oliver et al. 2004). The evidence to support this was limited to analyses of fasted/refed mouse livers for nascent rRNAs and by polysome profiles purporting to show reductions in 28S rRNA precursors and mature 60S subunits respectively. In contrast, RNAi depletion of RPL24 in human cell lines had no effect on ribosome biogenesis, or on the relative abundance of 40S and 60S subunits (Barkić et al. 2009; Wilson-Edell et al. 2014a). Furthermore, two reports have identified RPL24 as an exclusively cytoplasmic protein, leading to the hypothesis that it is assembled into mature ribosomes in the cytoplasm after rRNA synthesis and processing has already occurred (Barkić et al. 2009; Saveanu et al. 2003). In agreement with this, in the hindbrain of E9.5 embryos, the *Rpl24^Bst^* mutation had no effect on nucleolar architecture, indicating that it is not required for nucleolar function (Herrlinger et al. 2019). Our data agree with these later examples, as we fail to see any effect of the *Rpl24^Bst^* mutation on 60S:40S ratio in wild-type and transformed mouse intestines. Of further importance to the field, we also demonstrate in our tumour model that the translation defect in *Rpl24^Bst^* mutant mice is restored to normal levels following inactivation of eEF2K in *Rpl24^Bst^* mutant mice showing that the defect is dependent on eEF2K. From this, we conclude that ribosomes produced in *Rpl24^Bst/+^* mice would allow protein synthesis to proceed at near physiological levels, however this process is restricted via signalling through eEF2K/P-eEF2.

RPL24 depletion suppresses tumorigenesis in *Apc*-deficient *Kras*-mutant (APC KRAS) mouse models of CRC, but not in *Apc*-deficient *Kras*-wild-type (APC) models. In line with this, protein synthesis is reduced following depletion of RPL24 in our APC KRAS models, but not in APC models, despite induction of P-eEF2 in both cases. We previously showed that suppression of translation elongation via mTORC1/eEF2K/P-eEF2, using rapamycin, in the same APC models dramatically suppressed proliferation and extended survival (Faller et al. 2015). Thus, while RPL24 depletion or rapamycin treatment of *Apc*-deficient intestines each induce P-eEF2, only rapamycin treatment suppresses protein synthesis. The effect on protein synthesis appears to be the differential driving this divergence, with proliferation only impaired when protein synthesis is reduced. Crucially, RPL24 deficiency does not suppress mTORC1 activity since phosphorylation of the mTORC1 effectors 4E-BP1 and RPS6 (at S240/S244) are not reduced in *Apc*-deficient *Rpl24^Bst^*, providing an explanation of how proliferation and protein synthesis are maintained in *Apc*-deficient *Rpl24^Bst^* mutant animals. We also provide evidence that *Rpl24* mRNA is not elevated in line with other ribosomal protein mRNAs following *Kras*-mutation, which could explain why physiological expression levels of RPL24 are required for proliferation in *Kras*-mutant models.

It is interesting to reflect on previous work with the *Rpl24^Bst^* mutant mouse considering the mechanism we have described here. The *Rpl24^Bst^* mutation suppresses tumorigenesis in three mouse models of different types of blood cancer (Signer et al. 2014; Barna et al. 2008; Hsieh et al. 2010). In each case the *Rpl24^Bst^* mutation was found to decrease translation, although how the mechanism of translational suppression was not fully determined. Barna et al demonstrated a reduction in the cap-dependent translation using a luciferase reporter in *Rpl24^Bst^* MEFs (Barna et al. 2008). Furthermore, *Rpl24^Bst^* mutation dramatically suppressed cap-dependent translation following MYC activation, consistent with *Rpl24^Bst^* slowing tumorigenesis via reduced translation. We have shown that in mouse models of CRC the molecular mechanism by which normal levels of RPL24 maintains translation is via eEF2K and P-eEF2, therefore implicating this pathway in these previously studied blood cancer models.

The *Rpl24^Bst^* mutation has also been used to analyse brain development from neural progenitor cells, and explored as a model for retinal degenerative disease (Riazifar et al. 2015; Herrlinger et al. 2019). These studies found a revertant phenotype in neural progenitor cells overexpressing LIN28A and the defect in subretinal angiogenesis in *Rpl24^Bst/+^* mice. The role of translation elongation with respect to these phenotypes should now be analysed. eEF2K, and the many pathways it integrates, are potential targets for intervention in these *in vivo* models. It will also be of interest to determine how the inactivation of eEF2K affects the whole-body phenotypes of the *Rpl24^Bst^* mouse, such as the coat pigmentation and tail defects.

eEF2K has a pleiotropic effect in tumorigenesis, acting akin to a tumour suppressor or promoter dependent on the context (Knight et al. 2020b). In some cancers eEF2K promotes tumorigenesis. For example, under nutrient deprivation eEF2K acts as a pro-survival factor in transformed fibroblasts and tumour cell lines by suppressing protein synthesis to ensure survival (Leprivier et al. 2013). Here we show that eEF2K inactivation does not modify intestinal tumorigenesis. However, eEF2K is required for the suppression of tumorigenesis and protein synthesis following rapamycin/mTORC1 inhibition in APC cells (Faller et al. 2015) or *Rpl24^Bst^* mutation in APC KRAS cells (this work). Therefore, although eEF2K does not directly drive tumorigenesis, low eEF2K activity ensures that there is no blockade of translation or proliferation in tumour cells. Furthermore, eEF2K expression is required for drug or signalling responses that suppress tumorigenesis, giving it tumour suppressive activity. In accordance, CRC patients with low eEF2K protein expression suffer a significantly worse prognosis (Ng et al. 2019). Furthermore, we present mRNA and protein expression data showing that eEF2K is reduced in clinical CRC samples compared to normal tissue. This agrees with a model where low eEF2K allows rapid translation elongation to promote proliferation.

Using the same clinical data sets we demonstrate that both RPL24 and eEF2 are elevated in CRC, consistent with their roles in promoting translation and proliferation. This presents the possibility of direct targeting of either RPL24 or eEF2 for anti-cancer benefit. RNAi against RPL24 in human breast cancer cell lines dramatically reduced proliferation (Wilson-Edell et al. 2014b). Similarly, targeting eEF2 using a compound reported to slow its exit from the ribosome and thus the rate of protein synthesis, reduces cell line and patient derived organoid growth (Stickel et al. 2015; Keysar et al. 2020). These reports agree with the data presented here suggesting that targeting of RPL24 or eEF2 would be beneficial in CRC.

This work uncovers an unexpected role for the ribosomal protein RPL24 in the regulation of translation elongation, acting via eEF2K/P-eEF2 signalling. We demonstrate that depletion of RPL24 suppresses tumorigenesis in a pre-clinical mouse model of a solid tumour, supplementing the three reports of the mutation suppressing blood tumour models. We provide genetic evidence supported by molecular assays of translation elongation to demonstrate that RPL24 depletion activates eEF2K to elicit tumour suppression in our models. We also speculate as to the role of eEF2K in the previously published blood cancer models where RPL24 depletion was beneficial. This work provides additional evidence for the anti-tumorigenic role of the eEF2K in CRC, highlighting the potential for targeting translation elongation for this disease.

## Materials and methods

### Materials availability

The mouse strains used will be made available on request. However, this may require a Materials Transfer Agreement and/or a payment if there is potential for commercial application. We ourselves are limited by the terms of Materials Transfer Agreements agreed to by the suppliers of the mouse strains.

### Mouse studies

Experiments with mice were performed under a licence from the UK Home Office (licence numbers 60/4183 and 70/8646). All mice used were inbred C57BL/6J (Generation ≥8) and were housed in conventional cages with a 12-hour light/dark cycle and *ad libitum* access to diet and water. Mice were genotyped by Transnetyx in Cordova, Tennessee. Experiments were performed on mice between the ages of 6 and 12 weeks, both male and female mice were used. Sample sizes for all experiments were calculated using the G*Power software (Faul et al. 2009) and are shown in the figures or legends. Researchers were not blinded during experiments. The *Villin^CreER^* allele (El Marjou et al. 2004) was used for intestinal recombination by intraperitoneal (IP) injection of tamoxifen in corn oil at a final *in vivo* concentration of 80mg/kg. The *Apc* flox allele, *Apc^Min^*, *Kras^G12D^* lox-STOP-lox allele, *Eef2k^D273A^* and *Rpl24^Bst^* alleles were previously described (Jackson et al. 2001; Shibata et al. 1997; Gildish et al. 2012; Oliver et al. 2004; Moser et al. 1990). Tumour model experiments began with a single dose of tamoxifen after which mice were monitored until they showed signs of intestinal disease – paling feet from anaemia, weight loss and hunching behaviour. Tumours were scored macroscopically by counting the number visible after fixation of intestinal tissue. For short-term experiments, mice wild-type for *Kras* were induced on consecutive days (days 0 and 1) and sampled 4 days after the first induction (day 4). Mice bearing the *Kras^G12D^* allele were induced once and sampled on day 3 post-induction. Where indicated, 250μL of BrdU cell proliferation labelling reagent (Amersham Bioscience RPN201) was given by IP injection 2 hours prior to sampling. For the regeneration experiments, mice were exposed to 10 Gy of γ-irradiation at a rate of 0.423 Gy per minute from a caesium 137 source. They were then sampled 72 hours after irradiation.

### Histology and immunohistochemistry

Tissue was fixed in formalin and embedded in paraffin. Immunohistochemistry (IHC) staining was carried out as previously (Faller et al. 2015), using the following antibodies; BrdU (BD Biosciences #347580), P-eEF2 T56 (Cell Signaling Technology (CST) #2331), P-4E-BP1 T37/46 (CST #2855), P-RPS6 S235/6 (CST #4858), P-RPS6 S240/4 (CST #5364), RPL24 (Sigma Aldrich HPA051653) and Lysozyme (Dako A0099). The IHC protocol followed the Vector ABC kit (mouse #PK-6102, rabbit #PK-6101). RNAScope analysis was carried out according to the manufacturers guidelines (ACD) using probes to murine *Olfm4* (#311838). For all staining, a minimum of 3 biological replicates were stained and representative images used throughout. For BrdU scoring in short-term model experiments tissue was fixed in methanol:choloform:acetic acid at a ratio 4:2:1 then transferred to formalin and embedded paraffin.

### Intestinal organoid culture

Crypt cultures were isolated and then maintained as previously described (Knight et al, in press). In all crypt culture experiments, multiple biologically independent cultures were generated and analysed from different animals of the shown genotypes. DMEM/F12 medium (Life Technologies #12634-028) was supplemented with 5 mM HEPES (Life Technologies #15630-080), 100 U/mL penicillin/streptomycin (Life Technologies #1540-122), 2 mM L-glutamine (Life Technologies #25030-024), 1x N2, 1x B27 (Invitrogen #17502-048 and #12587-010), 100 ng/mL noggin (Peprotech #250-38) and 50 ng/mL EGF (Peprotech #AF-100-15). Wild-type cultures were also supplemented with 500 ng/mL R-spondin (R&D Systems #3474-RS). For *ex vivo* growth assays, cells were plated in technical triplicate in 96 well plates, and proliferation measured using Cell-Titer Blue (Promega #G8080) added to previously untreated cells each day for up to 4 days. Rapamycin (LC Laboratories R-5000) was dissolved in DMSO and administered for 24 hours, comparing to just DMSO vehicle treated cells.

### Western blotting

Samples were lysed (10 mM Tris (pH 7.5), 50 mM NaCl, 0.5% NP40, 0.5% SDS supplemented with protease inhibitor cocktail (Roche #11836153001) PhosSTOP (Roche #04906837001) and benzonase (Sigma Aldrich # E1014)) on ice and then the protein content estimated by BCA assay (Thermo Fisher Scientific #23225). 20μg of protein were denatured in loading dye containing SDS then resolved by 4-12% SDS-PAGE (Invitrogen #NP0336BOX). Protein was transferred to nitrocellulose membranes, blocked with excess protein and immunoblotted overnight at 4°C using the following antibodies; P-eEF2 T56 (CST #2331), eEF2 (CST #2332), RPL24 (Sigma Aldrich HPA051653) and β-actin (Sigma Aldrich #A2228) as a sample control. 1 hour incubation at room temperature with secondary antibodies (HRP-conjugated anti-mouse secondary (Dako #P0447); HRP-conjugated anti-rabbit secondary (Dako #P0448)) was followed by exposed to autoradiography films with ECL reagent (Thermo Fisher Scientific #32106). Quantification was performed using Image J (Rueden et al. 2017).

### Sucrose density gradients

Cells were replenished with fresh medium for the 6 hours before harvesting. This medium was then spiked with 200 µg/mL cycloheximide (Sigma Aldrich #C7695) 3 minutes prior to harvesting on ice. Crypt fractions from mice were isolated by extraction of the epithelial from 10cm of proximal small intestine. Each data point plotted represents an individual animal. PBS-flushed linearly-opened small intestines were incubated in RPMI 1640 medium (Thermo Fisher Scientific #21875059) supplemented with 10mM EDTA and 200µg/mL cycloheximide for 7 minutes at 37°C with regular agitation to extract villi. Crypts were isolated by transferring remaining tissue to ice-cold PBS containing 10mM EDTA and 200 µg/mL cycloheximide for a further 7 minutes, again with agitation. The remaining tissue was discarded. Samples were lysed (300 mM NaCl, 15 mM MgCl2, 15 mM Tris pH 7.5, 100 µg/mL cycloheximide, 0.1% Triton X-100, 2 mM DTT and 5 U/mL SUPERase.In RNase Inhibitor(Thermo Fisher Scientific #AM2696)) on ice and post-nuclear extracts placed on top of 10-50% weight/volume sucrose gradients containing the same buffer (apart from no Triton X-100, DTT, or SUPERase.In). These were then spun in an SW40Ti rotor at 255,000rcf for 2 hours at 4°C under a vacuum. Samples were then separated through a live 254 nM optical density reader (ISCO). Polysome to subpolysome (P:S), heavy to light polysome (H:L) or 60S to 40S (60S:40S) ratios were calculated using the manually defined trapezoid method. For harringtonine run-off assays, cultures were prepared in duplicate for each genotype. One was pretreated with 2µg/mL harringtonine (Sant Cruz #sc-204771) for 5 minutes (300 seconds) prior to cycloheximide addition. This was then processed as above. Run-off rates were calculated as previously (Knight et al. 2015).

### ^35^S methionine incorporation assay

Organoids were replenished with medium 6 hours prior to analysis while in the optimal growth phase post-splitting. Technical triplicates were used from 3 different animals per experiments. ^35^S-methionine (Perkin Elmer #NEG772002MC) was used at 30 µCi/mL for 30 minutes. Samples were prepared in technical triplicate then lysed using the same buffer described for western blotting. Protein was precipitated in 12.5% (w/v) trichloroacetic acid onto glass microfiber paper (Whatmann #1827-024) by use of a vacuum manifold. Precipitates were washed with 70% ethanol and acetone. Scintillation was read on a Wallac MicroBeta TriLux 1450 scintillation counter using Ecoscint scintillation fluid (SLS Ltd #LS271) from these microfiber papers. In parallel the total protein content was determined by the BCA assay using un-precipitated sample. Protein synthesis rate was expressed as the scintillation normalised to the total protein content (CPM / µg/mL protein), which was then changed to a relative value compared to relevant controls for each experiment.

### Publicly available clinical data analysis

The TNMplot and UALCAN web portals were used to analysis publicly available dataset from The Cancer Genome Atlas and Clinical Proteomic Tumor Analysis Consortium. Details of these portals are available in these publications (Bartha and Győrffy 2021; Chandrashekar et al. 2017).

### Statistical analyses

All statistical analyses are detailed in the relevant figure legends. In all cases, calculated *P* values less than or equal to 0.05 were considered significant. N numbers for each experiment are detailed within each figure, as individual points on graphs or within figure legends.

### Data availability

Source data for Figure 1F and 4F are available as part of this manuscript. All raw data used in this manuscript will be freely available on Dryad.

## Acknowledgements

The Sansom laboratory was funded by CRUK (A17196, A24388, A21139), The European Research Council ColonCan (311301). This work was also funded by a Wellcome Trust Collaborative Award in Science (201487) to GM, CMS, TvdH, AEW and OS. We are grateful to the Advanced Technologies and Core Services at the Beatson Institute (funded by CRUK C596/A17196 and A31287), particularly the Biological Services Unit, Histology Services and Transgenic Technology Laboratory. CGP is supported by funding from the National Health and Medical Research Council (Australia). We thank Fiona Warrander for critical reading of the manuscript.

## Author contributions

Study design: JRPK, OJS. Data acquisition: JRPK, NV, DMG, RAR, WJF. Data analysis: JRPK, NV. Data interpretation: JRPK, NV, DMG, CGP, GM, TvdH, CMS, AEW, OJS. Writing the manuscript: JRPK, AEW, OJS.

---

**Supplemental Figure 1:**
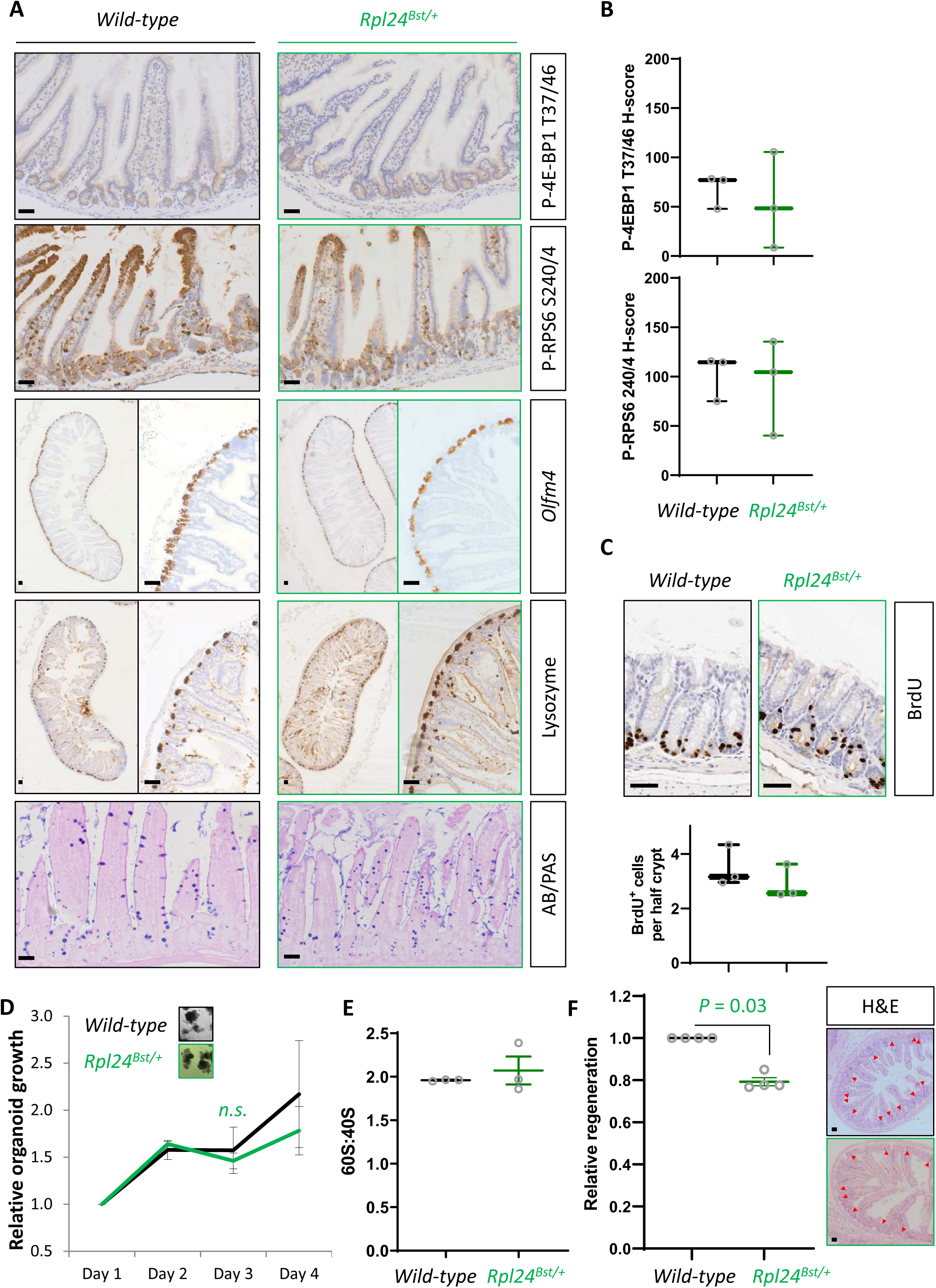
*Rpl24^Bst^* mutation does not affect intestinal homeostasis, or 40S:60S ratio: (A) Representative images of staining for intestinal lineages and translation associated phosphorylation sites – P-4E-BP1 T37/46, and P-RPS6 240/4 of intestinal sections from wild-type and *Rpl24^Bst/+^* mice. *Olfm4* defines stem cells, lysozyme for Paneth cells and AB/PAS (Alcian blue/periodic acid-Schiff) for goblet cells. Bars represent 50µm. (B) H-score quantification of P-RPS6 S240/4 and P-4E-BP1 T37/46 from the same experiment as in (A). (C) BrdU staining of the medial colon from wild-type and *Rpl24^Bst^* mice, top, with scoring of BrdU positive cells per half crypt below. Scores are from 3 mice per genotype, each plotted as the average of at least 20 half crypts. (D) Organoids either wild-type of mutant for *Rpl24^Bst^* were grown over 4 days and the change in growth plotted relative to day 1. Triplicate independent lines were used for each genotype, with the average of these plotted on the graph ±SEM. Lack of significance was determined by Mann Whitney U test. (E) Area under the curve for 40S and 60S ribosomal subunits was determined from the traces as in Figure 1E. Data are from three independent biological replicates ±SEM. (F) Wild-type or *Rpl24^Bst^* mice were given 10Gy of irradiation then sampled 72 h later. The number of regenerative crypts was quantified from at least 6 cross sections and the average plotted relative to wild-type regeneration set as 1 (n=4 per genotype). Micrograph insets show representative sections of intestine for wild-type (black box) and *Rpl24^Bst/+^* (green box) mice. Red arrows indicate regenerating crypts. Bars represent 50µm. Data are represented as the mean ±SEM with significance determined by Mann Whitney U test.

**Supplemental Figure 2:**
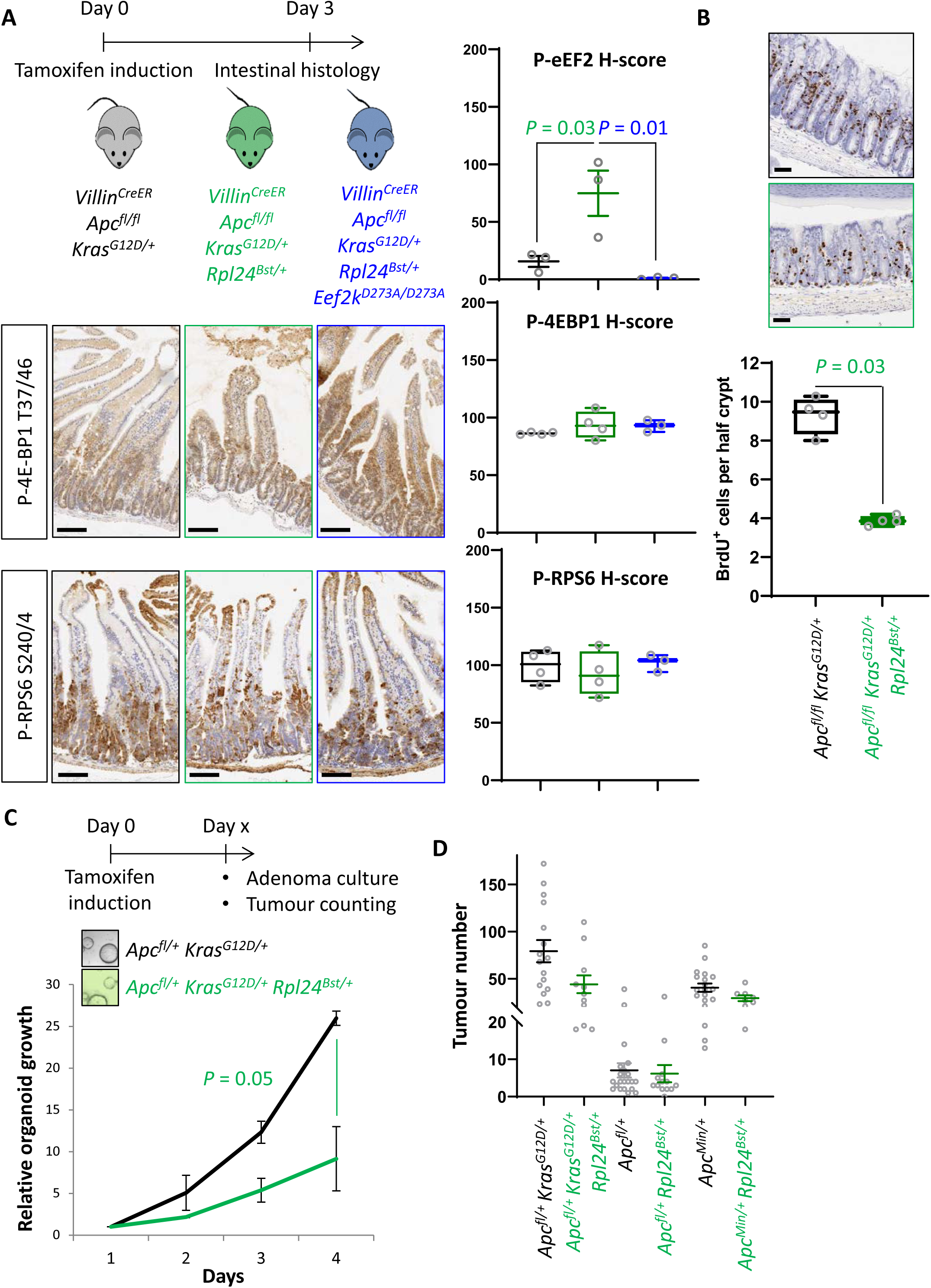
*Rpl24^Bst^* mutation leads to increased eEF2F phosphorylation and suppresses proliferation in colorectal cancer models. (A) Top: Schematic representation of experimental approach. *Villin^CreER^ Apc^fl/fl^ Kras^G12D/+^, Villin^CreER^ Apc^fl/fl^ Kras^G12D/+^ Rpl24^Bst/+^* or *Villin^CreER^ Apc^fl/fl^ Kras^G12D/+^ Rpl24^Bst/+^ Eef2k^D273A/D273A^* mice were induced by intraperitoneal injection of tamoxifen at 80mg/kg then intestinal tissue analysed 3 days later. Bottom: Staining for P-4E-BP1 T37/46 and P-RPS6 S240/4 from the genotypes above. Right: H-score quantification for P-eEF2 T56 (matched to images shown in Figures 2C and 5C), P-4E-BP1 T37/46 and P-RPS6 S240/4. Small intestines from at least 3 animals from each genotype (*Apc^fl/+^ Kras^G12D/+^*, *Apc^fl/+^ Kras^G12D/+^ Rpl24^Bst/+^* and *Apc^fl/fl^ Kras^G12D/+^ Rpl24^Bst/+^ Eef2k^D273A/D273A^*) were stained and the intensity quantified from the proliferative crypt region. Data are represented for 3 independent animals plotted ±SEM. Significance was determined by one-way ANOVA analysis with Tukey’s multiple comparison. (B) Top: Staining for BrdU in the medial colons of *Villin^CreER^ Apc^fl/fl^ Kras^G12D/+^* and *Villin^CreER^ Apc^fl/fl^ Kras^G12D/+^ Rpl24^Bst/+^* mice. Bottom: Representative micrograph images of each genotype. Scores are from 4 mice per genotype, each plotted as the average of at least 20 half crypts. Significance was determined by Mann Whitney U test. (C) Top: Schematic showing the generation of colonic adenoma cultures. of *Villin^CreER^ Apc^fl/+^ Kras^G12D/+^* and *Villin^CreER^ Apc^fl/+^ Kras^G12D/+^ Rpl24^Bst/+^* mice were induced and aged until colonic tumours were present. Individual adenoams were then excised and adenoma cells isolated. Bottom: Growth of these cultures over 4 days, plotted relative to day 1. Biologically independent triplicate organoid lines were used, with the averages plotted ±SEM. Significance was tested by Mann Whitney U test. (D) Tumour numbers for all tumour models reported, with and without *Rpl24^Bst^* mutation. Each point represents and individual mouse. From top to bottom n = 16, 11, 21, 13, 19 and 8 mice.

**Supplemental Figure 3:**
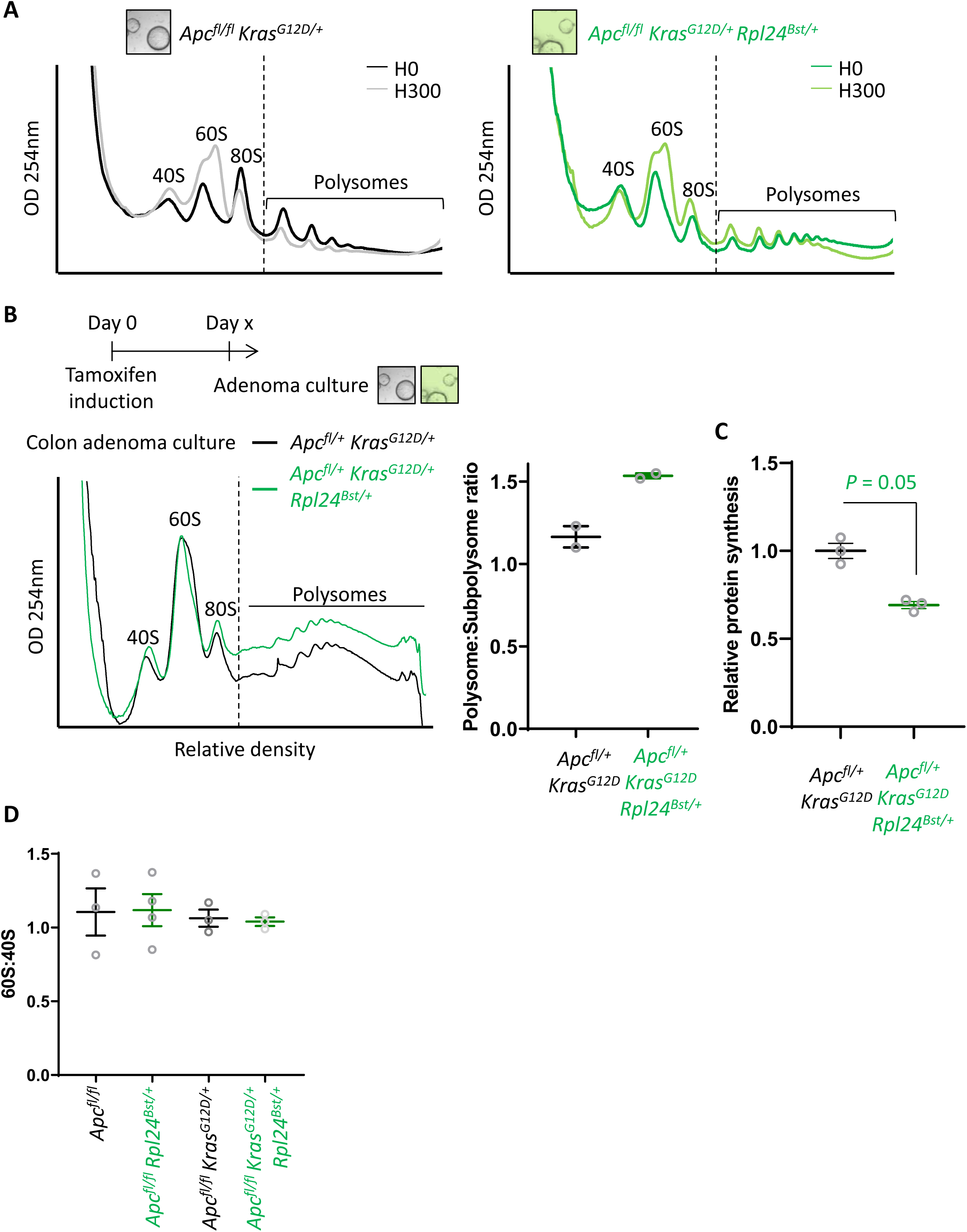
The effect of *Rpl24^Bst^* mutation on translation and ribosome composition: (A) Representative polysome profiles from *Apc^fl/fl^ Kras^G12D/+^* or *Apc^fl/fl^ Kras^G12D/+^ Rpl24^Bst/+^* small intestinal organoid cultures, pre-treated with harringtonine for 5 min / 300 s (H300) or untreated (H0). These traces were analysed for the run-off rates shown in Figure 3D. (B) Representative polysome profiles from *Apc^fl/+^ Kras^G12D/+^* or *Apc^fl/+^ Kras^G12D/+^ Rpl24^Bst/+^* colonic adenoma cultures (left) and quantification of the polysome to sub-polysome ratio from these (right). Two biologically independent lines were analysed per genotype and plotted ±SEM. Scheme above denotes the generation of adenoma cultures from distinct colonic tumours in aged *Villin^CreER^ Apc^fl/+^ Kras^G12D/+^* and *Villin^CreER^ Apc^fl/+^ Kras^G12D/+^ Rpl24^Bst/+^* mice. (C) Relative protein synthesis rates quantified by ^35^S methionine incorporation in the colonic adenoma cultures described in (B) with n=3. The average protein synthesis rates were plotted relative to *Apc^fl/+^ Kras^G12D/+^* controls (= 1) for 3 organoid lines per genotype ±SEM. Significance was determined by Mann Whitney U test. (D) 60S to 40S ratio from sucrose density gradients from lysates generated from the indicated genotypes. Data show the mean ±SEM. Representative traces are shown in Figure 3A and S4D.

**Supplemental Figure 4:**
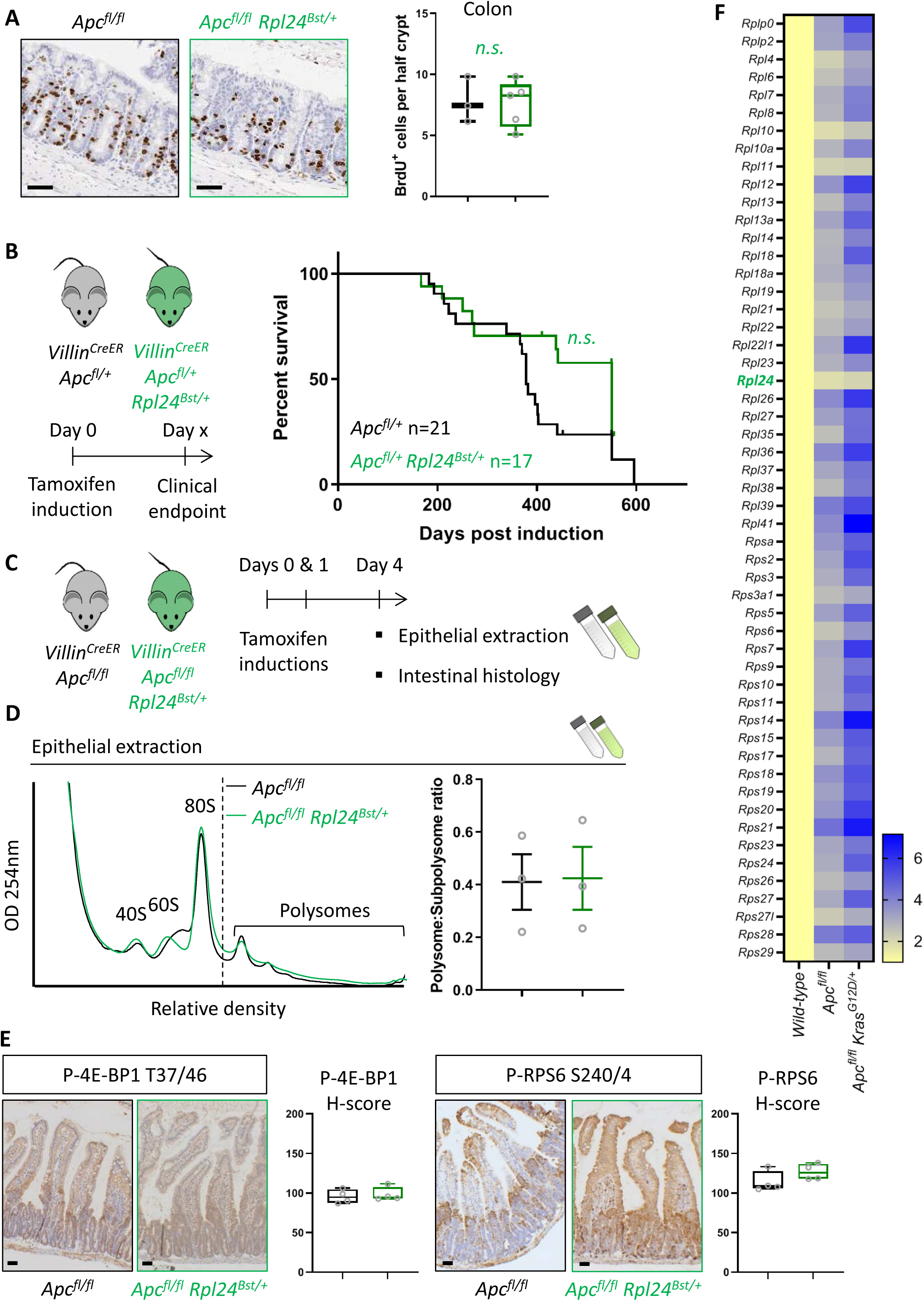
*Rpl24^Bst^* mutation has no benefit in models of CRC with wild-type *Kras*: (A) Left: Staining for BrdU in the medial colons of *Villin^CreER^ Apc^fl/fl^* and *Villin^CreER^ Apc^fl/fl^ Rpl24^Bst/+^* mice. Bottom: Scores are from 3 and 5 mice per genotype, each plotted as the average of at least 20 half crypts. Lack of significance was determined by Mann Whitney U test. (B) Left: Schematic of experiment. the *Villin^CreER^ Apc^fl/+^* mice with or without *Rpl24^Bst^* mutation were induced then aged until showing signs of intestinal tumours. Right: Survival curve from the *Villin^CreER^ Apc^fl/+^* tumour model, with and without *Rpl24^Bst^* mutation. Censored subjects were sampled for health reasons not relating to the intestine. Lack of a significant difference was determined by Mantel-Cox test. (C) Schematic representation of experimental approach. *Villin^CreER^ Apc^fl/fl^* or *Villin^CreER^ Apc^fl/fl^ Rpl24^Bst/+^* mice were induced by 2 intraperitoneal injection of tamoxifen at 80mg/kg on days 0 and 1 then intestinal tissue analysed on day 4. Intestines analysed histologically or by epithelial extraction for sucrose density gradient analysis. (D) Left: Representative polysome profiles generated from *Apc^fl/fl^* intestinal extracts with or without the *Rpl24^Bst^* mutation. Subpolysomal components and polysomes are labelled. Right: Quantification of the polysome:subpolysome ratio across 3 biologically independent replicates for each genotype. Data show the mean ±SEM. (E) Staining of small intestinal tissue for P-4E-BP1 T37/46 and P-RPS6 S240/4 from *Villin^CreER^ Apc^fl/fl^* or *Villin^CreER^ Apc^fl/fl^ Rpl24^Bst/+^* mice alongside H-score quantification from the proliferative zones of the intestines of 4 mice per genotype. (F) Relative ribosomal protein mRNA abundances from RNA sequencing of wild-type, *Villin^CreER^ Apc^fl/fl^* and *Villin^CreER^ Apc^fl/fl^ Kras^G12D/+^* whole small intestinal tissue. 3 independent biological samples were analysed per genotype with the averages used in this analysis. RNA sequencing reads for *Villin^CreER^ Apc^fl/fl^* and *Villin^CreER^ Apc^fl/fl^ Kras^G12D/+^* tissue was normalised to wild-type tissue set to 1. Values are shown horizontally scaled as fold changes to the wild-type tissue for each genotype.

**Supplemental Figure 5:**
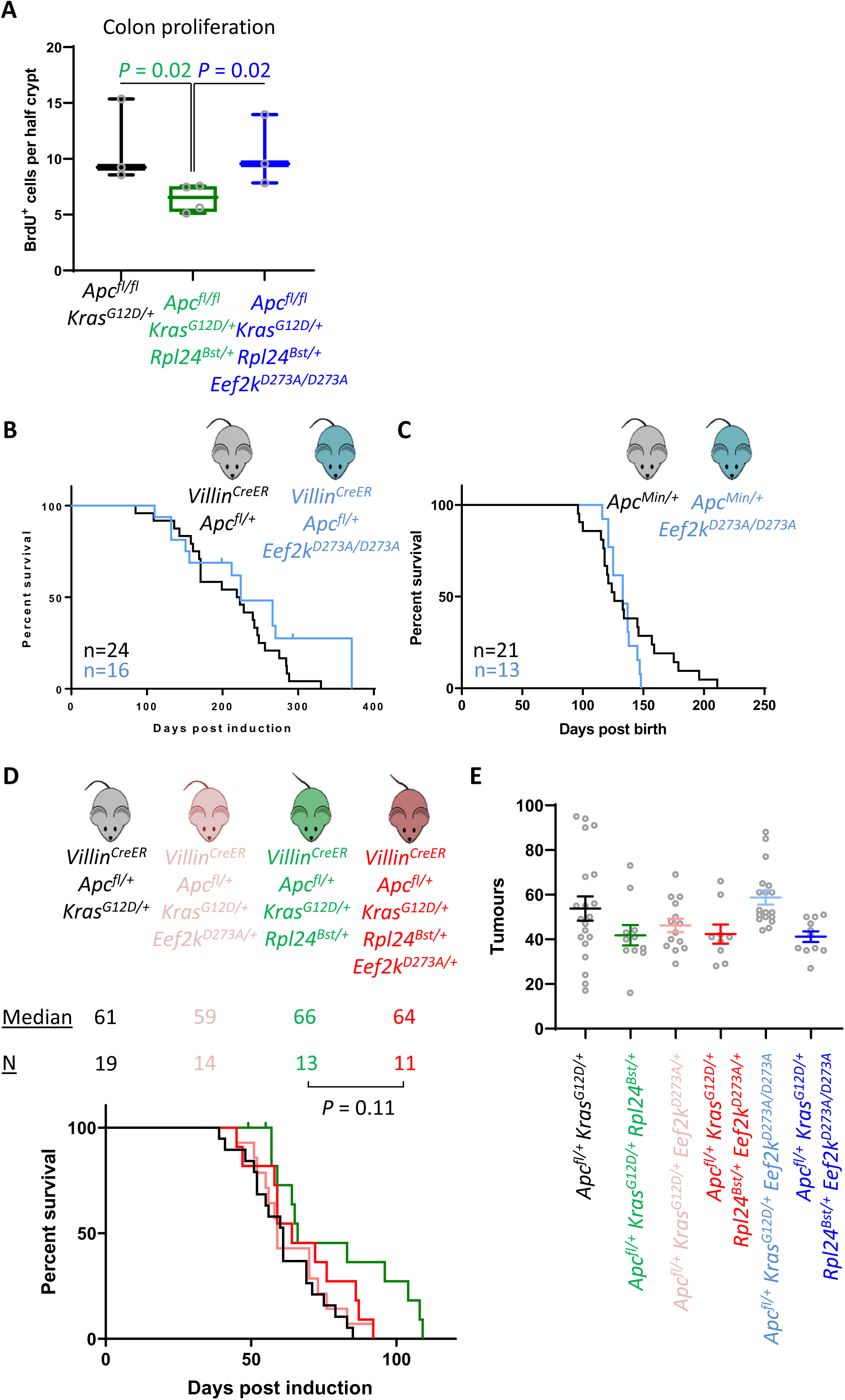
Heterozygous mutation of *Eef2k^D273A^*^/+^ partially suppresses the effects of *Rpl24^Bst^* mutation in the tumour model: (A) Scoring for BrdU incorporation in the medial colons of *Apc^fl/fl^ Kras^G12D/+^*, *Apc^fl/fl^ Kras^G12D/+^ Rpl24^Bst/+^* and *Apc^fl/fl^ Kras^G12D/+^ Rpl24^Bst/+^ Eef2k^D273A/D273A^* mice. Scores are from 3, 4 and 3 mice respectively per genotype, each plotted as the average of at least 20 half crypts. Significance was determined by one-way ANOVA analysis with Tukey’s multiple comparison. (B) Survival curve from the *Apc^fl/+^* tumour model for mice bearing the *Eef2k^D273A/D273A^* mutation or wild-type for *Eef2k*. Lack of a significant difference was determined by Mantel-Cox test. (C) *Apc^Min/+^* tumour model survival curve, for mice with wild-type *Eef2k* or the inactivating mutation, *Eef2k^D273A/D273A^*. Lack of a significant difference was determined by Mantel-Cox test. (D) Survival curve from the *Apc^fl/+^ Kras^G12D/+^* tumour model for mice bearing the *Rpl24^Bst^* mutation and/or one copy of the inactivating *Eef2k* mutation. Median survival in days and n numbers are shown for each genotype. Significance test was performed by Mantel-Cox test. *Apc^fl/+^ Kras^G12D/+^* and *Apc^fl/+^ Kras^G12D/+^ Rpl24^Bst/+^* survival curves are reused from Figure 5E. (E) Tumour numbers scored macroscopically from *Apc^fl/+^ Kras^G12D/+^* mice with or without *Rpl24^Bst^* mutation and with no, one or two copies of the inactivating *Eef2k^D273A^* mutation. Each point is an individual mouse with the bars depicting the mean ±SEM. From left to right n = 20, 11, 14, 9, 18 and 11 mice.

**Supplemental Figure 6:**
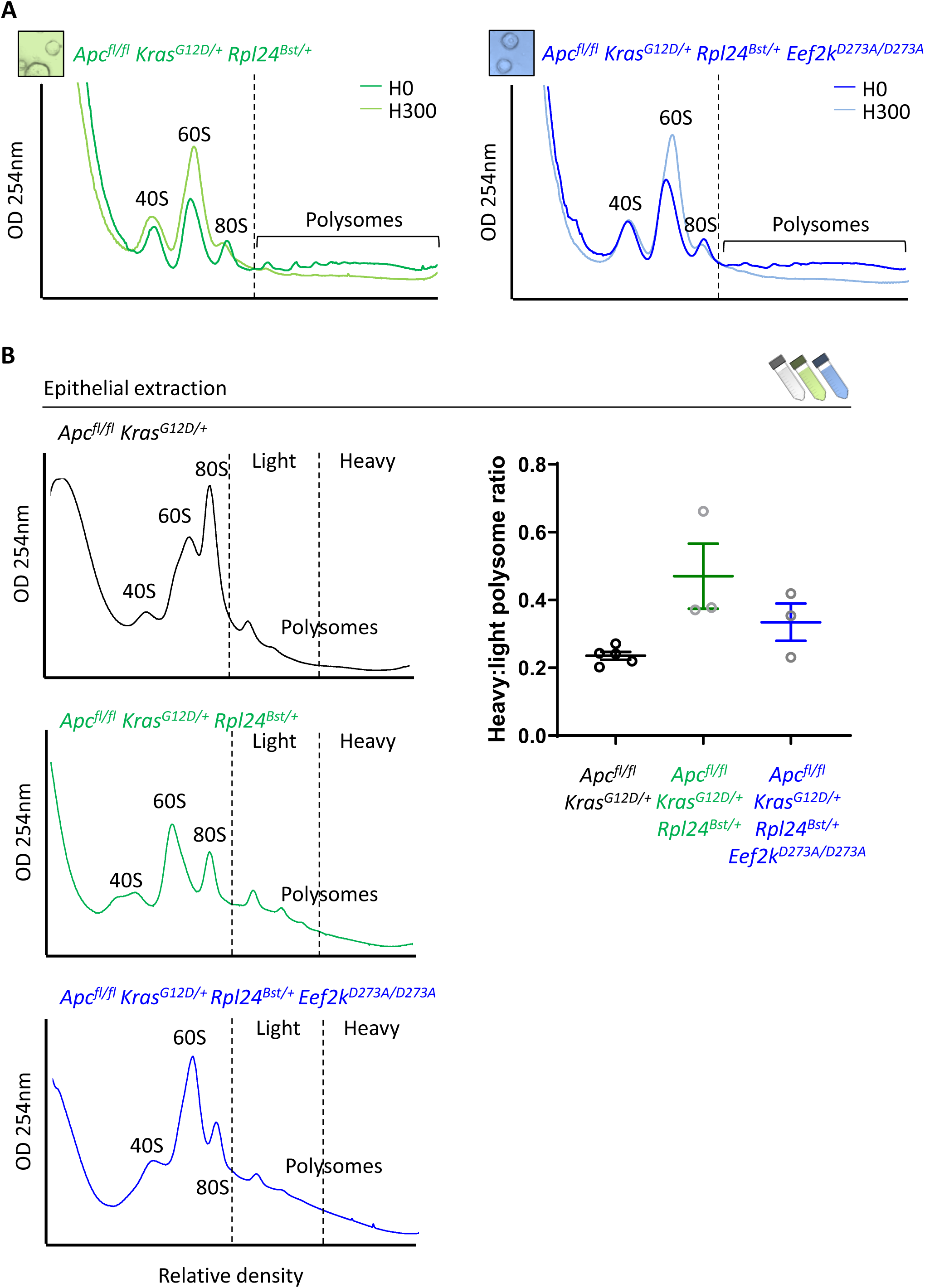
Inactivation of *Eef2k* restores translation elongation speed in *Rpl24^Bst^* mutant mice: (A) Polysome profiles from sucrose density gradients of *Apc^fl/fl^ Kras^G12D/+^ Rpl24^Bst/+^* or *Apc^fl/fl^ Kras^G12D/+^ Rpl24^Bst/+^ Eef2k^D273A/D273A^* small intestinal organoid cultures, pre-treated with harringtonine for 5 mins / 300 s (H300) or untreated (H0). These traces are representative of those analysed for the run-off rates shown in Figure 6B. (B) Representative sucrose density profiles generated from *Apc^fl/fl^ Kras^G12D/+^* intestinal extracts with or without the *Rpl24^Bst^* mutation and inactivation of eEF2K. Subpolysomal components and polysomes are labelled, and the polysomes have been split pictorially into light and heavy. To the right of this is quantification of the heavy:light polysome ratio. Data show the mean ±SEM of 5, 3 and 3 mice reading from left to right.

**Supplemental Figure 7:**
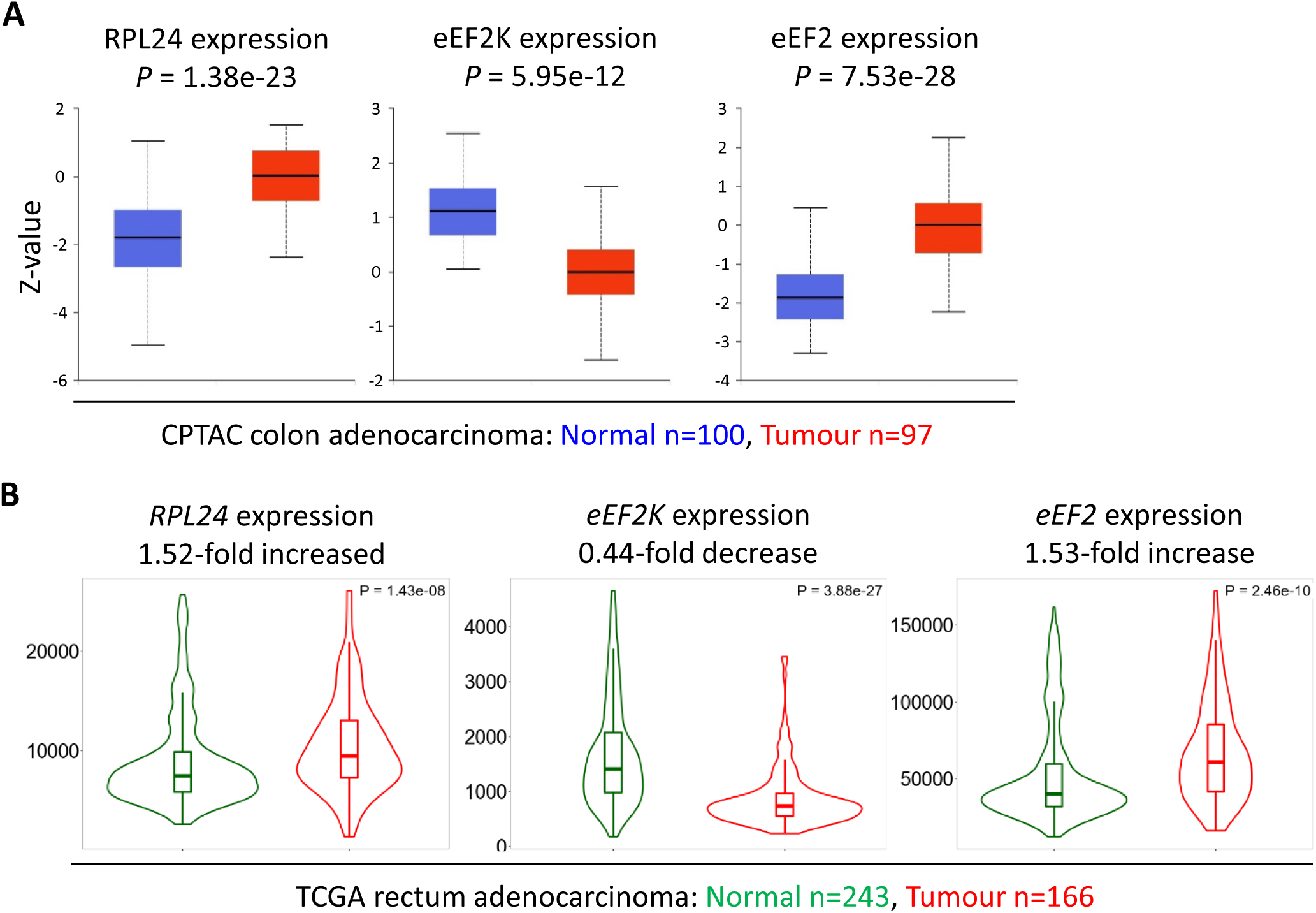
Expression of *RPL24*, *EEF2K* and *EEF2* suggest increased eEF2 activity in CRC tumours (A) Protein levels of RPL24, eEF2K and eEF2 in normal colon and colon adenocarcinoma collated from the Clinical Proteomic Tumor Analysis Consortium by UALCAN. Numbers of samples and *P* values are shown for each protein. (B) RNA expression levels of RPL24, EEF2K and EEF2 between normal rectum and rectum adenocarcinoma samples using data extracted from The Cancer Genome Atlas by TNMplot. Relative expression changes are annotated, as well as *P* values for each transcript.

## Notes

### Competing Interest Statement

The authors have declared no competing interest.

